# Disruption of DLL4/NOTCH1 Causes Dysregulated PPARγ/AKT Signaling in Pulmonary Arterial Hypertension

**DOI:** 10.1101/2024.01.31.578230

**Authors:** Keytam S. Awad, Shuibang Wang, Edward J. Dougherty, Ali Keshavarz, Cumhur Y. Demirkale, Zu Xi Yu, Latonia Miller, Jason M. Elinoff, Robert L. Danner

## Abstract

Pulmonary arterial hypertension (PAH) is a progressive cardiopulmonary disease characterized by vascular remodeling of small pulmonary arteries. Endothelial dysfunction in advanced PAH is associated with proliferation, apoptosis resistance, and endothelial to mesenchymal transition (EndoMT) due to aberrant signaling. DLL4, a cell membrane associated NOTCH ligand, activates NOTCH1 signaling and plays a pivotal role maintaining vascular integrity. Inhibition of DLL4 has been associated with the development of pulmonary hypertension, but the mechanism is incompletely understood. Here we report that *BMPR2* silencing in PAECs activated AKT and decreased DLL4 expression. DLL4 loss was also seen in lungs of patients with IPAH and HPAH. Over-expression of DLL4 in PAECs induced *BMPR2* promoter activity and exogenous DLL4 increased *BMPR2* mRNA through NOTCH1 activation. Furthermore, DLL4/NOTCH1 signaling blocked AKT activation, decreased proliferation and reversed EndoMT in *BMPR2*– silenced PAECs and ECs from IPAH patients. PPARγ, suppressed by BMPR2 loss, was induced and activated by DLL4/NOTCH1 signaling in both *BMPR2*-silenced and IPAH PAECs, reversing aberrant phenotypic changes, in part through AKT inhibition. Finally, leniolisib, a well-tolerated oral PI3K8/AKT inhibitor, decreased cell proliferation, induced apoptosis and reversed markers of EndoMT in *BMPR2*-silenced PAECs. Restoring DLL4/NOTCH1/PPARγ signaling and/or suppressing AKT activation may be beneficial in preventing or reversing the pathologic vascular remodeling of PAH.

## Introduction

Pulmonary arterial hypertension (PAH) is a progressive cardiopulmonary disease notable for vascular remodeling with obstruction and rarefaction of small pulmonary arteries (1). These functional and structural changes produce sustained increases in pulmonary vascular resistance, ultimately leading to right heart failure and death. Currently available therapies targeting the endothelin, nitric oxide and prostacyclin pathways have improve symptoms and prolong survival. However, stopping or reversing disease progression has remained elusive (2). Early endothelial injury and loss (3) followed by endothelial dysfunction with AKT activation (4) and a proliferative, anti-apoptotic phenotype are central hallmarks of PAH vascular pathology (5).

The PI3K/AKT signaling pathway plays an important role in cell survival and proliferation. In endothelial cells (ECs), the proliferative and anti-apoptotic activity of AKT can be regulated by the NOTCH pathway (6). This pathway consists of 4 transmembrane NOTCH receptors (NOTCH1-4) that interact with 5 membrane bound ligands (DLL1, DLL3, DLL4, JAGGED1 and JAGGED2) resulting in 2 cleavage events culminating in the release of the NOTCH intracellular domain (NICD) (7). NICD translocates to the nucleus where it heterodimerizes with the DNA binding protein RBP-J (recombination signal binding protein for immunoglobulin kappa J region) to regulate gene transcription. Highly conserved NOTCH signaling is an essential pathway that maintains vascular integrity by promoting endothelial cell cycle arrest (8), restricting endothelial inflammation (9) and stabilizing cell-to-cell junctions (10). Mice with targeted disruption of *Dll4* or *Notch1* die early in gestation of vascular defects.

Arterial endothelium predominantly expresses DLL4 (11), a NOTCH1 and NOTCH4 ligand. Notably, anti-DLL4 monoclonal antibodies, developed to treat cancer, have been associated with the development of pulmonary hypertension (PH) in clinical trials (12). Conversely, DLL4/NICD activation by olmesartan, an angiotensin II receptor blocker, attenuated transverse aortic constriction-induced cardiac remodeling in mice (13) and monocrotaline-induced pulmonary hypertension and right ventricular hypertrophy in rats (14). Importantly, BMPR2 was demonstrated to be required for NOTCH1 activation and deletion of endothelial specific *Notch1* in transgenic mice exacerbated hypoxia-induced pulmonary hypertension (15). Recently, Wang et al. demonstrated decreased NICD expression in PAH patients as well as in mouse models of pulmonary hypertension implicating NICD loss in the development of PAH (16).

While DLL4/NOTCH1 signaling appears to protect the pulmonary vasculature, how and under what conditions DLL4 loss or inhibition contributes to the development of pulmonary hypertension and pathologic vascular remodeling is not understood. Since BMPR2 (17) and PPARγ (18) both guard against the development of PAH (19, 20), we hypothesized that loss of DLL4/NOTCH signaling might result in pulmonary hypertension through interactions with BMPR2 and PPARγ. Here, BMPR2, DLL4/NOTCH and PPARγ cross-talk and interdependent regulation were investigated in healthy and *BMPR2*-silenced PAECs, as well as in lung from both IPAH and heritable PAH (HPAH) patients. These relationships were linked to effects on signal transduction pathways including AKT and JNK and to endothelial cell phenotypic changes associated with PAH pathogenesis.

## Methods

*Immunofluorescence.* Explanted lung tissue from unused donor controls, IPAH and HPAH subjects were obtained from PHBI (Table E1 for donor information). Frozen lung sections were air dried, fixed with fresh 4% paraformaldehyde for 10 min, washed 3x with PBS at room temperature (RT) for 10 min and permeabilized with 0.02% Trition X-100 for 5 min. Tissue sections were briefly dipped in PBS, blocked with 10% donkey serum for 1 h in the dark with gentle shaking and incubated with anti-CD31 and anti-pAKT (S473) overnight at 4° C. Sections were gently washed 3x with PBS, incubated with Alexa fluor 488 donkey anti-mouse and Alexa fluor 647 donkey anti-rabbit for 1 h at RT, washed 3x with PBS and stained with Hoechst 33342 for 15 min. Tissue sections were mounted with Prolong Diamond Antifade Mountant (ThermoFisher Scientific, Rockford, IL) and scanned using the Zeiss Axioscan Z1 (Zeiss, White Plains, NY).

*Immunohistochemistry.* Explanted human lung tissue from unused donor controls, IPAH and HPAH subjects were obtained from PHBI (Table E2 for donor information). Paraffin embedded lung tissue sections (10 μm) were cut (Histoserv, Inc., Germantown, MD) and slides were stained for expression of DLL4 at the Lombardi Comprehensive Cancer Center Histopathology and Tissue Shared Resource (Georgetown University Medical Center).

*Statistics.* Differences of relevant gene expression, protein expression/phosphorylation, caspase activity, apoptotic cells, cell migration, and BrdU incorporation between control and *BMPR2* siRNA transfected PAECs were analyzed by paired t-test. Differences between failed donor control and IPAH ECs were examined by unpaired t-test. ANOVA followed by post hoc Tukey’s honest significance test was used to analyze effects of *BMPR2*, *JNK1* or *CAV1* silencing and/or the efficacy of pathway inhibitors in PAEC apoptosis, proliferation, migration, gene expression and protein phosphorylation assays. Dose-responses across multiple conditions and donors were subjected to linear mixed models with random subject effects to account for the correlation within each donor. Log-transformation was applied when necessary. Two tailed *P* < 0.05 was accepted as significant. All statistical analyses were performed using JMP® software version 16.1.0 (SAS Institute Inc., Cary, NC).

## Results

*Silencing BMPR2 Activates AKT and Blocks Apoptosis.* Heterozygous loss-of-function (LOF) mutations in *BMPR2* are the leading cause of HPAH and accounts for 10-40% of cases without a family history (21). Aberrant activation of the AKT pathway has been associated with endothelial and smooth muscle cell proliferation as well as apoptosis resistance in late-stage PAH(4, 22). Therefore, the effect of BMPR2 loss on phosphorylation of both AKT and BCL2-associated agonist of cell death (BAD) were examined in PAECs after 48 h of knockdown (Figure 1A). *BMPR2* silencing significantly activated AKT as shown by phosphorylation of both S473 and T308, while BAD, a target of AKT, was phosphorylated at S136 which inactivates its pro-apoptotic effects. Compared to control lung tissue, immunofluorescence staining of small pulmonary arteries in lung tissue from PAH patients showed increased phosphorylation of AKT (S473) co-localized with CD31, an endothelial cell marker (Figure 1B; for patient characteristics see Table E1). Activation of AKT was also increased in plexogenic lesions as compared to control (Figure 1C). Caspase 3/7 activity, a marker of apoptosis, was suppressed in *BMPR2*-silenced PAECs grown in complete media and also after the induction of apoptosis by serum and growth factor withdrawal (Figure 1D).

**Figure 1.**
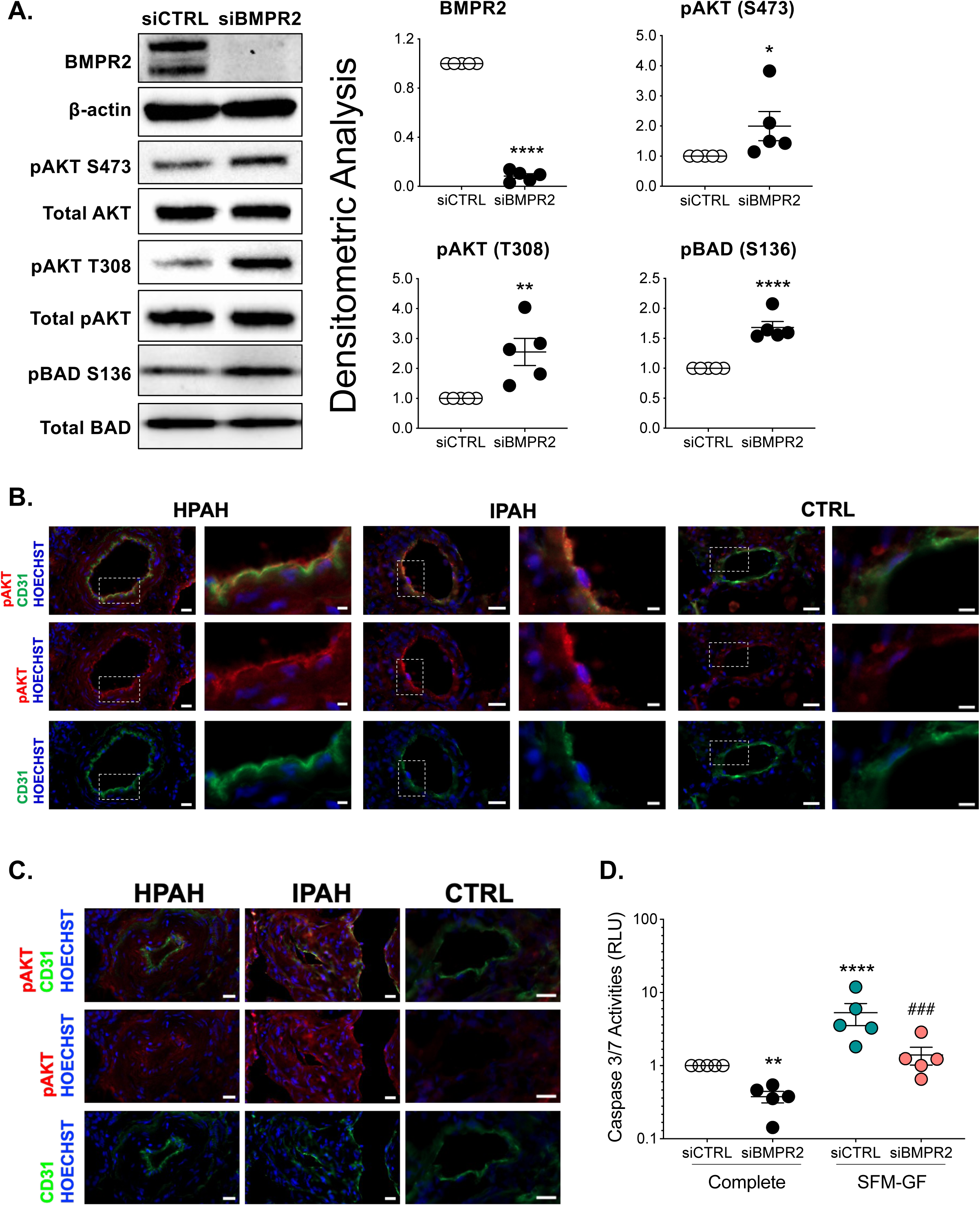
BMPR2 knockdown activates AKT and protects the cells against apoptosis. **(A)** Human primary pulmonary artery endothelial cells were transfected with control (siCTRL) or BMPR2 siRNA (siBMPR2) for 48 h and total protein lysates were collected. Western blots of BMPR2, phosphorylation of AKT at S473 and T308, total AKT, phosphorylation of BCL2-associated agonist of cell death (BAD) at S136 and total BAD were detected. Densitometric analysis of each protein relative to total β-actin, total AKT or total BAD and normalized to its corresponding siCTRL. Representative Western blots are shown (n=5). **(B-C)** Immunofluorescence of pAKT (S473) (red), CD31 (green) antibodies and nuclear staining with Hoechst 33342 (blue) in HPAH, IPAH and control (CTRL) lung. Scale bar, 20 μm. Insert scale bar, 5 μm. **(D)** Caspase 3/7 activity was measured after 24 h withdrawal of serum and growth media (SFM-GF; n=5). Data presented as mean ± SEM; (A) paired t-test, \**P* < 0.05; \*\**P* < 0.01; \*\*\*\**P* < 0.001; (C) 2-way ANOVA with Tukey HSD **^###^***P* < 0.005 (siBMPR2-complete vs siBMPR2-SFM-GF)

*JNK1 Suppression Contributes to Apoptosis-resistance.* We previously observed that BMPR2 loss in PAECs decreased JNK1 phosphorylation (23) a known downstream consequence of AKT activation (24). As JNK1 activation can trigger apoptosis, its suppression was examined as a possible contributor to the apoptosis resistant endothelial phenotype. *BMPR2* or *JNK1* silencing (Figure 2A) each similarly reduced both cleaved caspase 3, a measure of caspase 3 activity (Figure 2B), and proapoptotic BIM protein (Bcl-2 interacting mediator of cell death or BCL2L11; Figure 2C), while also increasing ERK phosphorylation/activation, another cell survival signal (Figure 2D). Moreover, flow cytometry analysis of annexin V and propidium iodide (AV/PI) staining of apoptotic cells showed that JNK1 knockdown, like BMPR2 loss, similarly protected cells from serum and growth factor withdrawal-induced apoptosis (Figure 2E and 2F).

**Figure 2.**
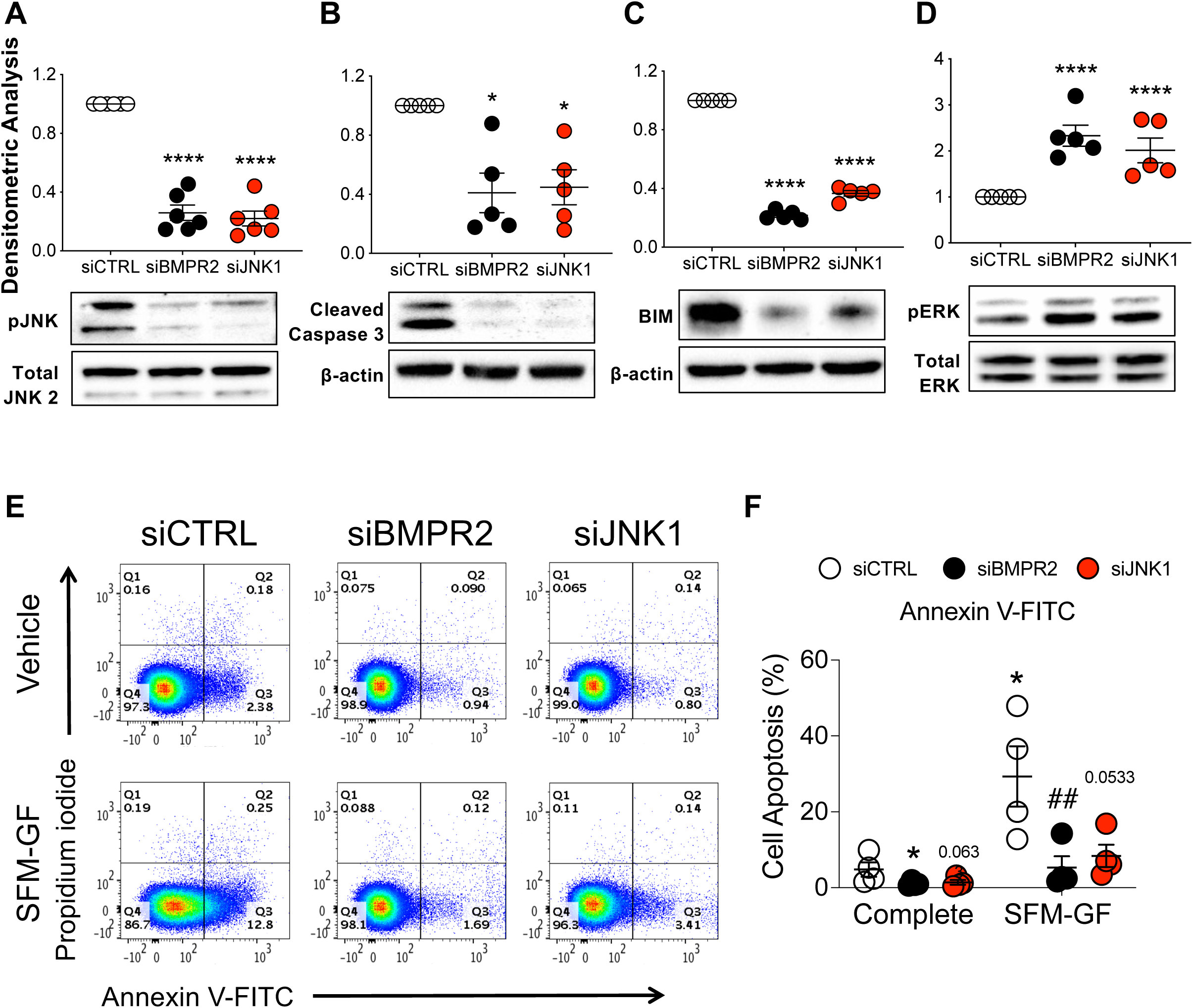
Loss of activated JNK1 contributes to apoptosis resistance as measured by reduced caspase 3 cleavage and flow cytometry analysis. Human primary pulmonary artery endothelial cells (PAECs) were transfected with control (siCTRL), BMPR2 (siBMPR2), or JNK1 (siJNK1) siRNA for 48 h and total protein lysates were collected. **(A)** phosphorylated JNK1 (pJNK), **(B)** cleaved caspase 3, **(C)** BIM and **(D)** phosphorylated ERK (pERK) protein levels were measured and densitometric analysis was performed relative to total JNK2, β-actin or total ERK and normalized to its corresponding siCTRL. Representative Western blots are shown (n=5). **(E)** After siRNA transfection (48 h), cells were cultured in either complete media or serum-free media without growth factors (SFM-GF) for 24 h followed by staining with annexin and propidium iodide (PI) for 15 min and data acquired on MACSquant. **(F)** Column scatter plot representing the percentage of apoptotic cells (cells stained with annexin V not PI, quadrant 3). Data presented as mean ± SEM; (A-D) 1-way ANOVA with Tukey HSD, \**P* < 0.05; \*\*\*\**P* < 0.001; (F) 2-way ANOVA with Tukey HSD; ^##^*P* < 0.01 (siCTRL-SFM-GF vs siBMPR2-SFM-GF)

*BMPR2 or JNK1 Silencing Both Blocks DLL4-mediated NOTCH Signaling.* Besides its role in apoptosis, JNK1 signaling contributes to lung microvascular angiogenesis and network formation (25), essential functions in the vasculature that overlap with NOTCH signaling (26). Notably, inhibition of JNK1 activity has been reported to decrease DLL4 (27) and NOTCH1 (28) expression, indicating that AKT suppression of JNK1 signaling in our PAEC model of PAH might also regulate DLL4 and therefore NOTCH signaling. Here, *BMPR2* or *JNK1* silencing in PAECs reduced DLL4 protein (Figure 3A) and N1ICD (Figure 3B), the transcriptionally active, intracellular effector of NOTCH1 signaling. Nonetheless, total NOTCH1 was unchanged (Figure 3C), while N2ICD was increased (Figure 3D). N4ICD was modestly reduced in *JNK1*-silenced PAECs and unchanged in *BMPR2*-silenced PAECs (Figure 3E). Note that N3ICD was not examined because NOTCH3 is not expressed in endothelium.

**Figure 3.**
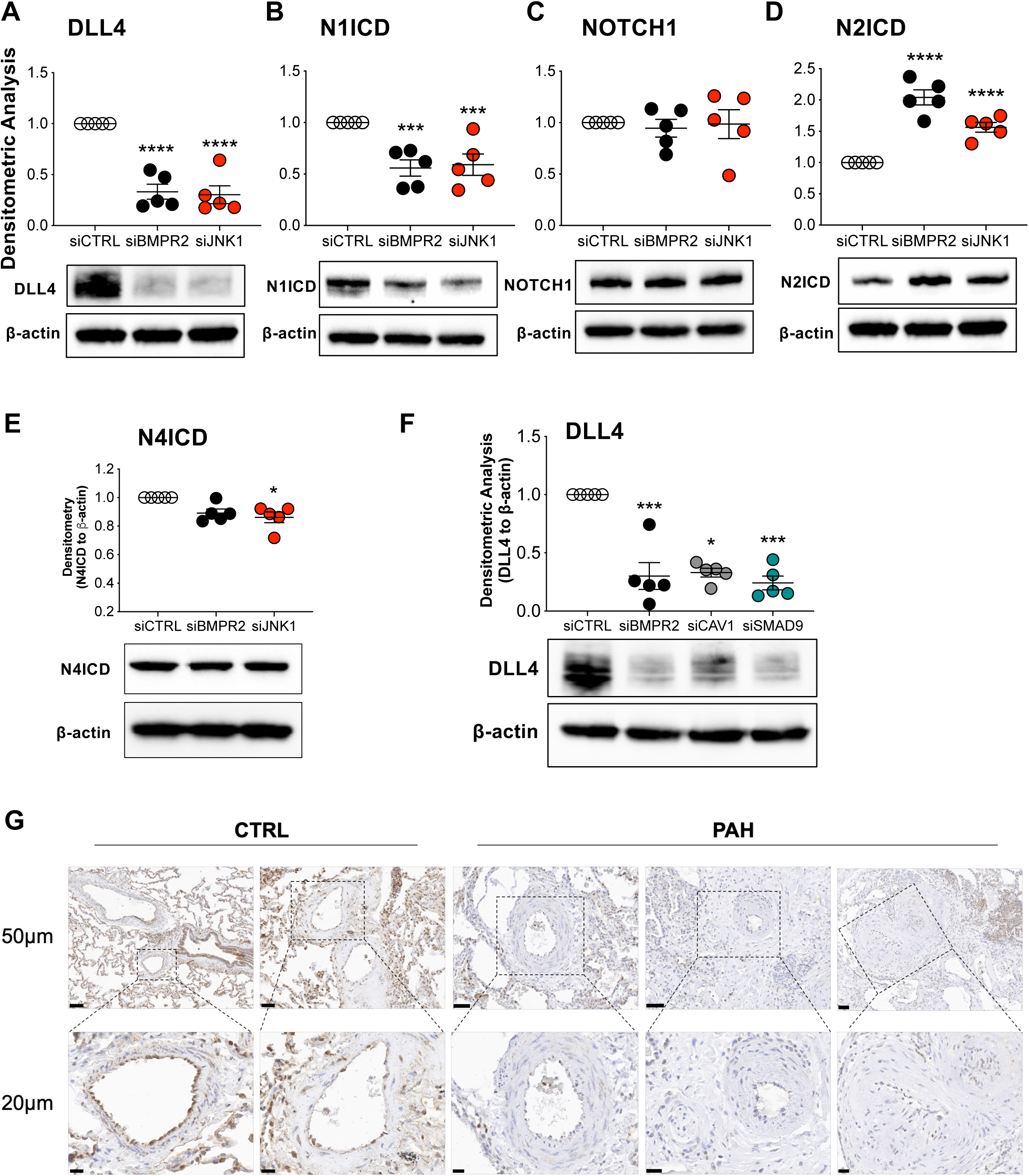
DLL4, a target of JNK1, is decreased in BMPR2– or JNK1-silenced PAECs and in lung tissue from patients with pulmonary arterial hypertension (PAH). Human primary pulmonary artery endothelial cells (PAECs) were transfected with control (siCTRL), BMPR2 (siBMPR2), or JNK1 (siJNK1) siRNA for 48 h and total protein lysates were collected for Western blotting of **(A)** DLL4, **(B)** N1ICD, **(C)** NOTCH1, **(D)** N2ICD and **(E)** N4ICD. **(F)** DLL4 protein was also analyzed in PAECs transfected with either control (siCTRL) or BMPR2 (siBMPR2), CAV1 (siCAV1) or SMAD9 (siSMAD9) gene-specific siRNA pools. CAV1 and SMAD9 loss-of-function mutations, like BMPR2, have been associated with the development of PAH. Densitometric analysis relative to β-actin and normalized to its corresponding siCTRL. Representative Western blots are shown (n=5). **(G)** Immunohistochemical staining of DLL4 in paraffin embedded lung of failed donor controls (CTRL; n=5), HPAH (n=3) and IPAH (n=2). Scale bar, 50μm and 20μm. Data are presented as mean ± SEM; 1-way ANOVA with Tukey HSD \**P* < 0.05; \*\*\**P* < 0.005; \*\*\*\**P* < 0.001

DLL4 expression was also examined in two other *in vitro* models of PAH, our *CAV1* (29) and *SMAD8/9* LOF models, both of which have been shown to activate AKT (unpublished data for SMAD8/9). Like *BMPR2* and *JNK1* silencing, *CAV1* and *SMAD8/9* loss similarly decreased DLL4 expression in human PAECs (Figure 3F). Notably, compared to healthy lung vessels, DLL4 protein expression was reduced in the small pulmonary artery endothelium of PAH lungs (Figure 3G and Figure E1; for patient characteristics see Table E2).

*Immobilized DLL4 Induces Endothelial N1ICD, Activates the BMPR2 Promoter and Increases BMPR2 mRNA.* To investigate if DLL4 can activate NOTCH signaling in cultured ECs, PAECs were grown in plates coated with either BSA or DLL4. Compared to BSA, immobilized DLL4 markedly activated NOTCH1 signaling as indicated by the accumulation of N1ICD and loss of NOTCH1 in both the absence and presence of *BMPR2* silencing (Figure 4A). N4ICD was not affected, indicating that DLL4-mediated NOTCH signaling was occurring primarily through NOTCH1 and not the NOTCH4 receptor. Although immobilized DLL4 blocked BMPR2 loss-induced increases in N2ICD (Figure 4A), this is unlikely to mediate the DLL4-driven EC phenotypes as NOTCH2/N2ICD suppression promotes proliferation and inhibits apoptosis in ECs (30, 31).

**Figure 4.**
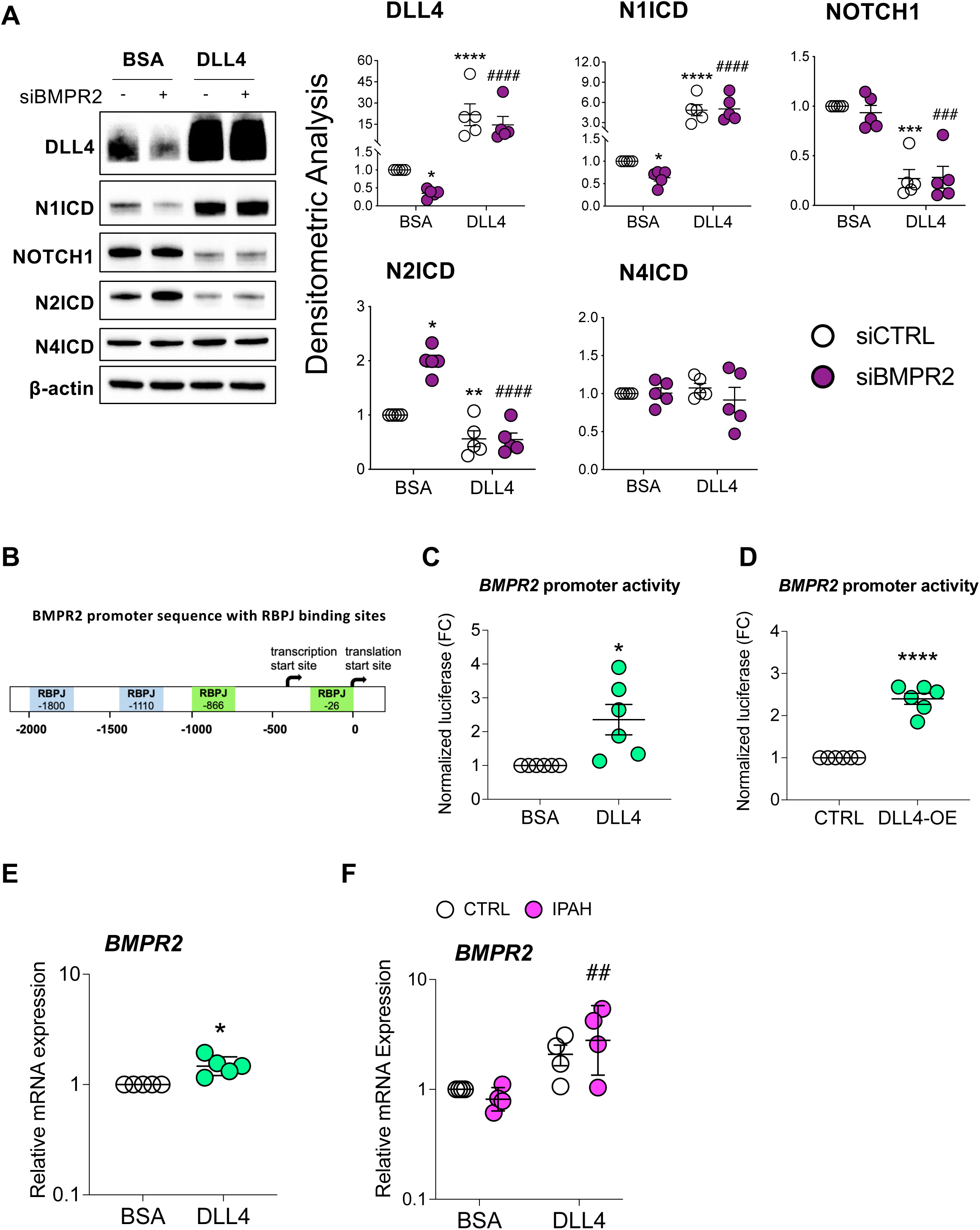
Immobilized DLL4 activates NOTCH1 signaling and increases *BMPR2* transcription. (**A**) Human primary pulmonary artery endothelial cells (PAECs) were grown on BSA or DLL4-coated plates and the following day transfected with either non-targeting control (siCTRL) or BMPR2 (siBMPR2) gene-specific siRNA pools. After BMPR2 knockdown (48 h), total protein lysates were collected and analyzed by Western blotting for expression of DLL4, N1ICD, NOTCH1, N2ICD and N4ICD. Representative Western blots are shown (n=5). Densitometric analysis (mean±SEM) relative to β-actin and normalized to its corresponding siCTRL-BSA. **(B)** The schematic of *BMPR2* promoter with putative RBPJ (recombinant signal binding protein) binding sites indicated. **(C)** BMPR2 promoter activity in PAECs grown on BSA or DLL4 coated plates and transfected with the promoter driven luciferase reporter. **(D)** BMPR2 promoter activity in PAECs transfected with the promoter driven luciferase reporter plus either control (CTRL) or DLL4 over-expression plasmid (DLL4-OE). **(E** and **F)** Quantitative RT-PCR of *BMPR2* mRNA from **(E)** PAECs or from **(F)** healthy or IPAH PAECs grown on BSA or DLL4 coated plates. For quantitative RT-PCR, data are presented as the geometric mean±SD (n=5). 2-way ANOVA with Tukey HSD; \**P* < 0.05; \*\**P* < 0.01; \*\*\**P* < 0.005; \*\*\*\**P* < 0.001 (siCTRL-BSA versus siBMPR2-BSA); **^##^***P* < 0.01; **^###^***P* < 0.005; **^####^***P* < 0.001 (siBMPR2-BSA versus siBMPR2-DLL4)

Despite the reports that loss of N1ICD in endothelium can cause PH (16), the precise mechanism is not known. Interestingly, TRANSFAC^®^ analysis identified two binding motifs for a Notch pathway transcription factor (recombination signal binding protein for immunoglobulin kappa J region; RBPJ) within the *BMPR2* promoter (Figure 4B), suggesting that NOTCH signaling may directly regulate *BMPR2* transcription. In support of this, PAECs grown on immobilized DLL4 (Figure 4C) or transfected with a *DLL4* overexpression plasmid (Figure 4D) each activated a reporter gene driven by the *BMPR2* promoter. DLL4 over-expression and N1ICD activation was verified in PAECs (Figure E2). Moreover, *BMPR2* mRNA was increased in PAECs grown on DLL4 coated plates (Figure 4E) regardless of whether these cells were from failed donor controls or IPAH patients (Figure 4F).

*DLL4-induced Endothelial N1ICD Signaling Blocks AKT Activation and Suppresses Cell Proliferation and EndoMT.* Next, we investigated whether immobilized DLL4-induced NOTCH1 signaling could block BMPR2 loss-associated AKT and ERK activation. *BMPR2* silencing increased AKT phosphorylation at both S473 and T308 which was suppressed by DLL4-induced NOTCH1 signaling (Figure 5A). The same was true for ERK activation. Because constitutive AKT activation has been demonstrated to induce proliferation and markers of EndoMT, we examined whether DLL4/NOTCH1 signaling affected cell proliferation or EndoMT. As previously reported (23), *BMPR2* silencing increased while DLL4/NOTCH1 activation inhibited PAEC proliferation (Figure 5B). Cell proliferation was also compared between failed donor control (CTRL) and IPAH PAECs. Although no difference was detected between these two groups when cultured *ex vivo* on BSA coated plates, exogenous DLL4 significantly inhibited proliferation in both (Figure 5C). In PAECs from IPAH patients compared to failed donor controls, expression of α-SMA (ACTA2), a mesenchymal marker, was significantly increased and this effect was significantly abrogated by immobilized DLL4 as assessed by immunofluorescence staining (Figure 5D). Conversely, VE-cadherin (CDH5), an EC marker, was decreased in IPAH compared to failed donor cells and was modestly enhanced by DLL4, but only in the IPAH ECs (Figure 5E). Furthermore, *BMPR2*-silenced PAECs or IPAH PAECs grown on DLL4 demonstrated reduced α-SMA protein as compared to BSA (Figure E3A and E3B). Together, these data correspond to current notions of PAH pathogenesis including increased PAEC proliferation and EndoMT and suggest that DLL4-induced NOTCH1 signaling plays a critical role in the regulation of both BMPR2 and AKT. This may provide a mechanistic explanation for the development of pulmonary hypertension in patients treated with monoclonal antibodies that block DLL4.

**Figure 5.**
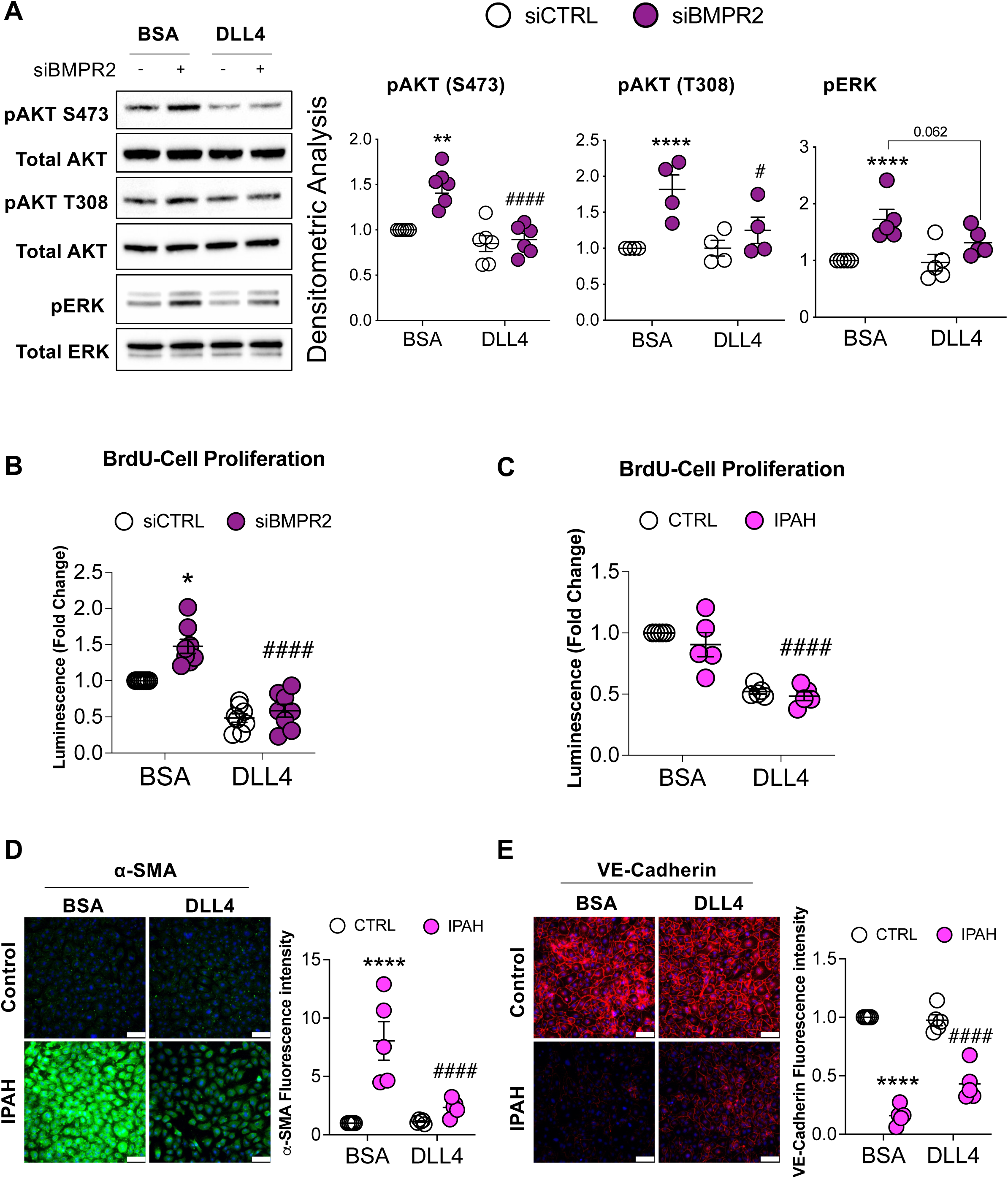
DLL4-induced NOTCH1 activation blocks AKT activation, inhibits proliferation and reverses EndoMT. (**A**) Human primary pulmonary artery endothelial cells (PAECs) were grown on BSA or DLL4-coated plates and the following day, cells were transfected with either non-targeting control (siCTRL) or BMPR2 (siBMPR2) gene-specific siRNA pools. After BMPR2 knockdown (48 h), total protein lysates were collected and analyzed by Western blotting for expression of phosphorylated AKT (pAKT S473 and pAKT T308) or phosphorylated ERK (pERK). Representative Western blots are shown (n=4-6, as indicated). Densitometric analysis (mean ± SEM) relative to total AKT or total ERK and normalized to its corresponding siCTRL. **(B)** BrdU cell proliferation of PAECs transfected with control or BMPR2 siRNA for 48 h and then replated onto BSA or DLL4-coated plates for an additional 48 h (n=5). **(C)** BrdU cell proliferation of healthy and IPAH ECs were grown on either BSA or DLL4 for 48 h (n=5). **(D** and **E)** Immunofluorescence staining of **(D)** α-SMA (ACTA2) and **(E)** VE-Cadherin (CDH5) in ECs from healthy and IPAH grown on either BSA or DLL4. Scale bar, 100μm. Data are presented as mean±SEM, 2-way ANOVA with Tukey HSD; \**P* < 0.05; \*\*\*\**P* < 0.001 (siCTRL-BSA versus siBMPR2-BSA); **^#^***P* < 0.05; **^##^***P* < 0.01; **^###^***P* < 0.005; **^####^***P* < 0.001 (siBMPR2-BSA versus siBMPR2-DLL4)

*DLL4/N1ICD Upregulation of PPARγ Expression and Signaling Reduces AKT Activation*. Loss of PPARγ in ECs is characterized by a proliferative, apoptosis-resistant phenotype (20) similar to the phenotype seen following loss of BMPR2. PPARγ is also a downstream effector of BMPR2 (19), therefore we next examined whether DLL4-mediated NOTCH1 signaling affected PPARγ expression. Notably, immunofluorescence staining for PPARγ protein in PAECs from IPAH patients was decreased, while DLL4-induced N1ICD restored PPARγ expression in these cells (Figure 6A). Furthermore, PAECs transfected with a *DLL4* overexpression plasmid and a PPAR-driven reporter showed that DLL4/N1ICD signaling increased the transcription of PPARγ target genes (Figure 6B). As expected, transfection of PAECs with the *DLL4* overexpression plasmid increased *DLL4* mRNA levels similar to exposure with immobilized DLL4, and was also found to increase *PPARG* mRNA (Figure E4A and E4B). Accordingly, in PAECs from IPAH patients (Figure 6C) and in *BMPR2*-silenced PAECs (Figure 6D), DLL4 not only induced *PPARG* transcription, but also increased mRNA expression of several PPARγ target genes (*FABP4*, *CYP1A1*, *PGK1* and *HK1*). Finally, because immobilized DLL4 induced PPARγ signaling while blocking AKT activation, the ability of PPARγ to suppress AKT phosphorylation and therefore activation was examined in human PAECs. Over-expression of PPARγ was found to decrease AKT phosphorylation at T308 (Figure 6E), implicating PPARγ as a mechanistic link between DLL4-mediated NOTCH1 signaling and AKT inhibition.

**Figure 6.**
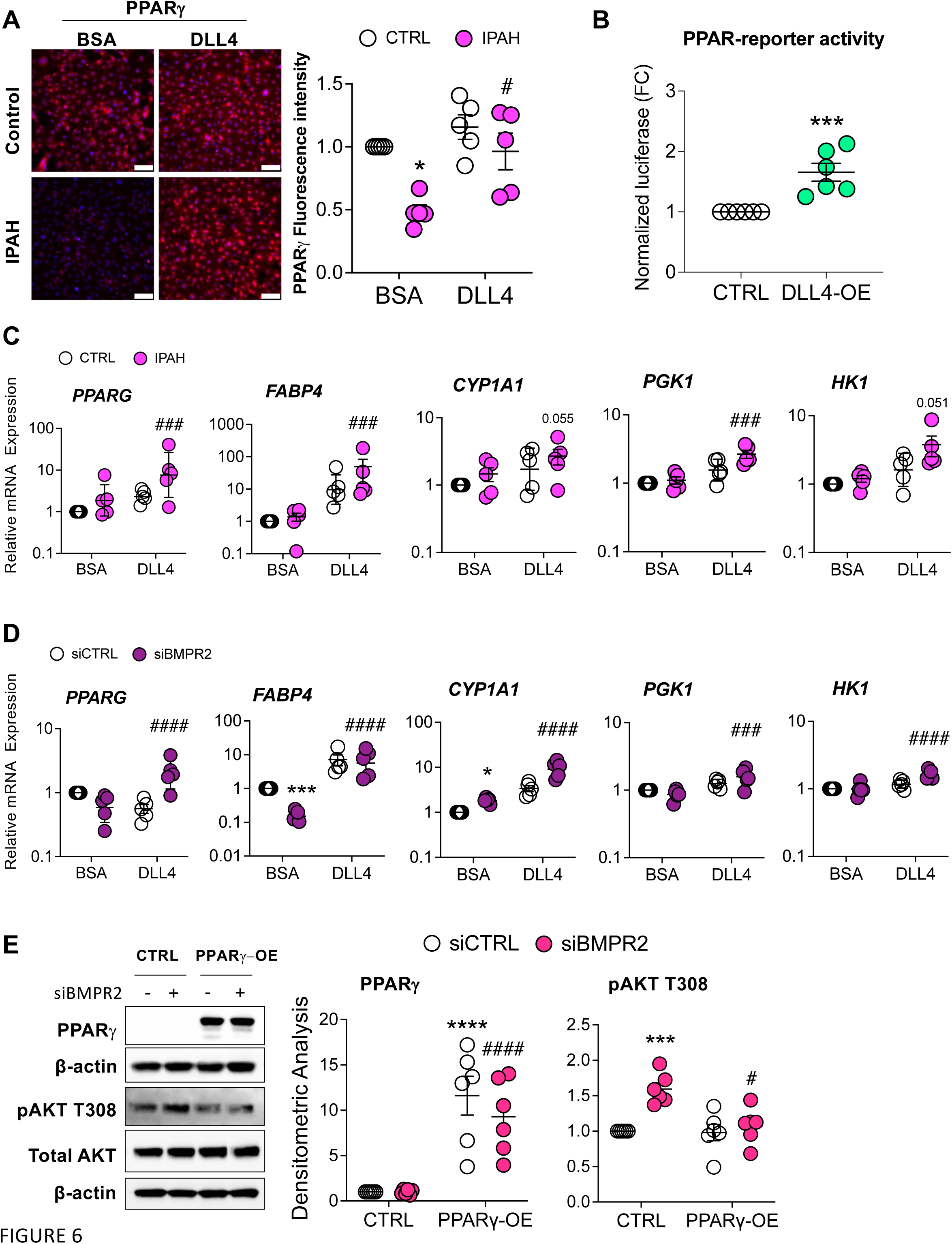
DLL4-induced NOTCH1 activation rescues PPARγ expression. (**A**) PPARγ immunofluorescence staining of PAECs endothelial cells from healthy or IPAH patients grown on either BSA or DLL4 (n=5). Scale bar, 100 μm. **(B)** Human primary pulmonary artery endothelial cells (PAECs) were transfected with empty vector (CTRL) or DLL4 overexpression plasmid (DLL4-OE) for 24 h and PPAR-driven reporter activity assessed (n=5). **(C)** mRNA levels from healthy (CTRL) or IPAH ECs grown on BSA or DLL4 for 48 h were analyzed for *PPARγ* expression and PPARγ target genes. **(D)** mRNA from PAECs transfected with control (siCTRL) or BMPR2 (siBMPR2) siRNA and grown on BSA or DLL4 for 48 h were analyzed for *PPARγ* expression and PPARγ target genes. **(E)** After 24 h of gene silencing with either control or BMPR2 siRNAs, PAECs were then transfected with empty vector (CTRL) or PPARγ overexpression plasmid for 24 h and expression of PPARγ and phosphorylated AKT (pAKT T308) protein was analyzed by Western blotting. Densitometric quantification (mean±SEM) relative to β-actin or total AKT and normalized to its corresponding siControl. Representative Western blots are shown (n=5 or 6). The mRNA levels were measured by quantitative RT-PCR and presented as the geometric mean ± SD (n=5). 2-way ANOVA with Tukey HSD; \**P* < 0.05; \*\*\**P* < 0.005 (siCTRL-BSA versus siBMPR2-BSA); **^#^***P* < 0.05; **^###^***P* < 0.005; **^####^***P* < 0.001 (siBMPR2-BSA versus siBMPR2-DLL4)

*Leniolisib Attenuates AKT Activation, Inhibits Cell Proliferation and EndoMT, and Induces Apoptosis in BMPR2 and CAV1 Silenced PAECs.* AKT activation and signaling have been implicated in PAH pathogenesis through its pro-survival, anti-apoptotic actions (4, 22). Because DLL4/NOTCH1 signaling increased BMPR2/PPARγ expression, suppressed AKT activation and reversed cell proliferation and EndoMT, PI3K/AKT was targeted to determine whether it would similarly ameliorate the aberrant phenotypic changes associated with BMPR2 loss. TNFα, an inflammatory cytokine associated with PAH progression (32), has been reported to specifically induce PI3Kδ in ECs (33). Because of its recent FDA approval for the treatment of children with activated phosphoinositide 3-kinase delta (PI3Kδ) syndrome (APDS) and tolerability in that population (34, 35), leniolisib was tested for its ability to block BMPR2 loss-induced AKT activation. First, TNFα was confirmed to induce PI3Kδ and activate AKT phosphorylation at S473 in PAECs (Figure E5A). Importantly, TNFα stimulation similarly increased PI3Kδ in both control and *BMPR2*-silenced PAECs (Figure E5B). Next, leniolisib was found to dose-dependently decrease BMPR2 loss-induced AKT activation at S473 (Figure 7A) in human PAECs. In addition to *BMPR2* silencing, leniolisib also mitigated AKT activation at both S473 and T308 in our *CAV1* LOF model (29) of PAH (Figure 7B).

**Figure 7.**
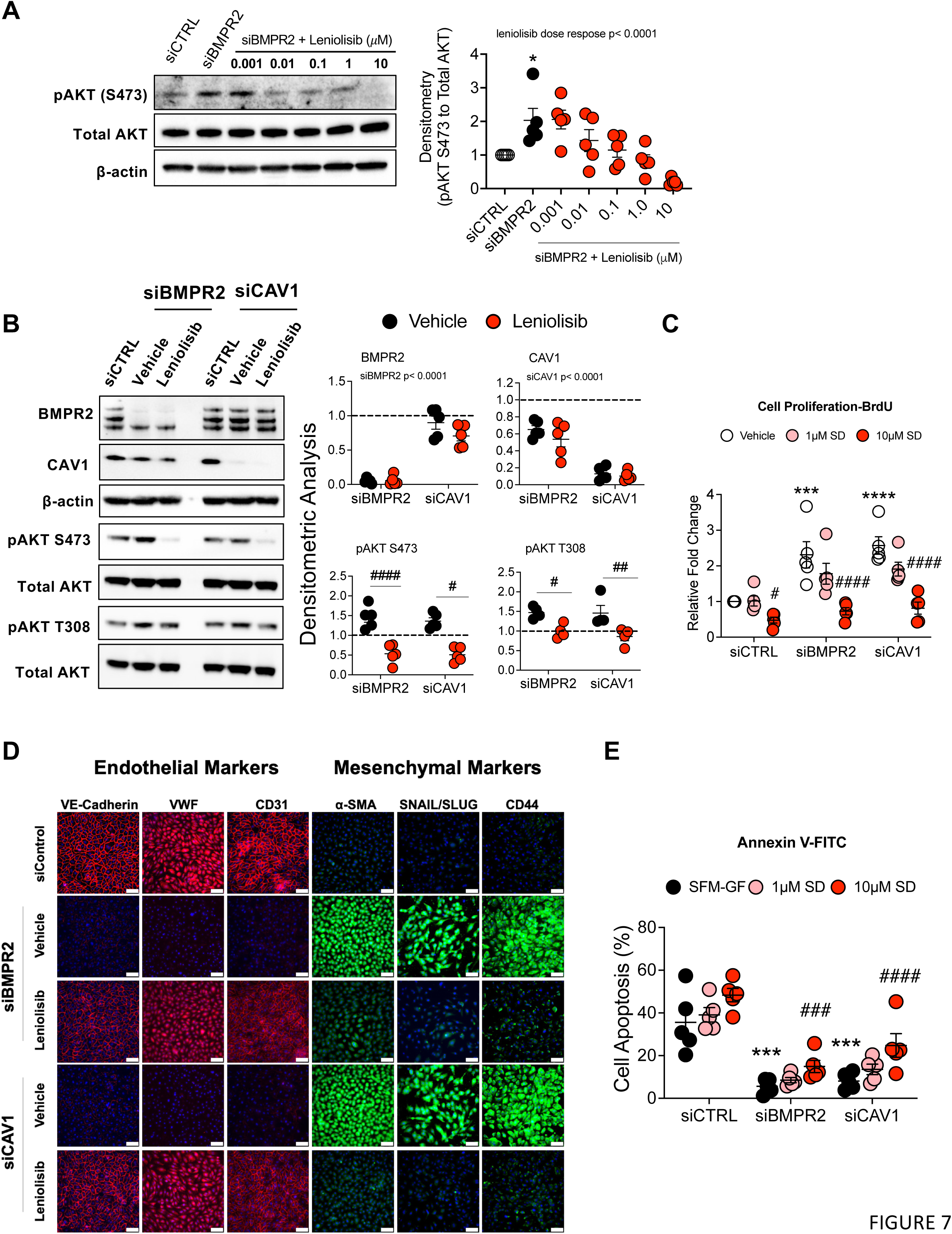
PI3K8 inhibitor, leniolisib, decreases AKT activation, proliferation, and EndoMT while increasing apoptosis in *BMPR2*-silenced PAECs. (**A**) Human primary pulmonary artery endothelial cells (PAECs) were transfected with control (siCTRL) or BMPR2 (siBMPR2) siRNA for 48 h and then exposed to leniolisib for 4 h at the indicated concentrations. **(B)** Effect of 10 μM leniolisib on phosphorylation of AKT (S473 and T308) in *BMPR2*– and *CAV1*-silenced PAECs. Representative Western blots are shown (n=5) and densitometric analysis (mean±SEM) relative to β-actin or total AKT and normalized to its corresponding siCTRL. **(C)** BrdU cell proliferation of PAECs transfected with control (siCTRL), BMPR2 (siBMPR2) or CAV1 (siCAV1) siRNA for 48 h then replated and treated with vehicle (DMSO), 1 μM or 10 μM leniolisib for an additional 72 h. **(D)** Representative immunofluorescence of endothelial markers (VE-cadherin, VWF and CD31) and mesenchymal markers (α-SMA, SNAIL/SLUG and CD44) in PAECs transfected with control, BMPR2, or CAV1 siRNA for 48 h then treated with 10 μM leniolisib for 24 h (n=5). Scale bar, 100 μm. **(E)** Leniolisib reactivated apoptosis as assessed by annexin and PI staining. \**P* < 0.05 (siCTRL versus siBMPR2), # *P* < 0.05 (vehicle versus treatment)

Finally, we determined the effect of leniolisib on cell proliferation, EndoMT and apoptosis, all of which have been associated with the endothelial dysfunction of PAH (5). Increased cell proliferation in both *BMPR2* and *CAV1* LOF models (23, 29), was markedly reduced by leniolisib at concentrations that were achievable in patients with activated PI3Kδ syndrome (35) (Figure 7C). Forty-eight hours after *BMPR2* or *CAV1* silencing, PAECs exhibited reduced expression of endothelial markers (VE-cadherin, VWF and CD31) and enhanced expression of mesenchymal markers (α-SMA, SNAIL/SLUG and CD44) as assessed by immunofluorescence. All of these changes were significantly reversed by leniolisib (Figure 7D; fluorescence intensity of each EndoMT marker was quantified and plotted in Figure E6). Importantly, leniolisib dose-dependently increased apoptosis as measured by AV/PI staining (Figure 7E) and enhanced caspase 3/7 activity in two of our *in vitro* LOF models of PAH (Figure E7).

## Discussion

Mutations that inactivate *BMPR2* are the most frequent genetic defects associated with the development of PAH (36) and reduced BMPR2 expression is also common in IPAH (37). Endothelial dysfunction manifested by proliferation, apoptosis resistance and EndoMT contributes to small pulmonary artery remodeling, a hallmark of late-stage PAH progression (5). Here, AKT activation was implicated in this dysfunctional phenotype. DLL4/NOTCH1 signaling was identified as a regulator of AKT, as well as an essential inducer of both *BMPR2* and *PPARG* in human PAECs. Likewise, BMPR2 loss itself was found to disrupt DLL4/NOTCH1 signaling, suppressing PPARγ expression and activating AKT. While BMPR2 deficiency reduced DLL4 protein and DLL4/NOTCH1 signaling, two other *in vitro* models of PAH based on *CAV1* or *SMAD8/9* LOF were also associated with reduced DLL4 expression. More importantly, small pulmonary arteries from patients with IPAH and HPAH both exhibited elevated AKT activation and reduced DLL4 expression compared to vessels from donor lungs. Accordingly, exogenous or overexpressed DLL4 both suppressed AKT activation, while also inducing itself, *BMPR2*, and *PPARG* in PAECs, including those from IPAH patients. Likewise, PPARγ overexpression was also found to block AKT activation in *BMPR2*-silenced PAECs. Finally, exogenous DLL4 or leniolisib, a PI3Kδ/AKT inhibitor, were shown to prevent the aberrant PAH-like endothelial phenotype associated with BMPR2 loss. These novel findings demonstrate extensive crosstalk between the DLL4/NOTCH1 and BMPR2/PPARγ networks in the regulation of AKT, supporting the development of strategies to activate DLL4/NOTCH1/PPARγ and/or inhibit AKT signaling for the treatment of pathologic vascular remodeling in PAH.

LOF or haploinsufficiency of *DLL4* (38) or *NOTCH1* (39) lead to Adams-Oliver syndrome, which includes PAH as one of its manifestations. DLL4 monoclonal antibodies, which were associated with PH in cancer patients (12), have been recently shown to cause pulmonary vascular remodeling and PH in mice through the inhibition of endothelial NOTCH1 signaling (16). Interestingly, *SOX17* deficiency, a risk factor for the development of IPAH, HPAH and PAH associated with congenital heart disease (40), was reported to transcriptionally induce DLL4 and protect against vascular leakage (41). In fact, loss of endothelial specific *SOX17* (42) in mice mimicked *DLL4* haploinsuffiency (43) suggesting that *SOX17* mutations might cause PAH in part by disrupting DLL4 signaling. However, this hypothesis has not been tested. These previous studies from different groups are consistent with our results and support the conclusion that impaired DLL4/NOTCH1 signaling plays a role in PAH pathogenesis. Other reports not supporting this conclusion warrant mention. For example, in human PAECs and umbilical endothelial cells, γ-secretase inhibitors, which block the proteolytic cleavage and release of intracellular N1ICD from the NOTCH1 receptor, were demonstrated to decrease cell proliferation and survival, right ventricular systolic pressure, and right heart hypertrophy in sugen/hypoxia rats (44, 45). These discrepant results might be due to the nonselectivity of γ-secretase inhibitors, targeting other ligand/NOTCH pairs and not just DLL4/NOTCH1. Additionally, γ-secretase may regulate other receptors, such that γ-secretase inhibitors might have non-NOTCH, off-target effects. Notably, DLL4 levels were not measured, and exogenous DLL4 was not used in these investigations.

JNK1, a downstream target of AKT (24) is a multi-functional protein that regulates important physiological processes such as apoptosis, proliferation and angiogenesis. AKT antagonizes JNK1 kinase activity by inhibiting upstream kinases including apoptosis signal-regulating kinase 1 (ASK1), MKK4/MKK7 and MLK (24). In mice, loss of JNK1 signaling in ECs was reported to suppress Dll4/Notch signaling (27). Likewise, Jnk1 KO mice were found to have reduced NOTCH1 expression (28). These findings suggest that AKT activation, followed by downstream JNK1 inactivation, could promote and reinforce the inhibition of DLL4 mediated NOTCH signaling. Indeed, our present study showed that AKT was activated and JNK was suppressed in *BMPR2*-silenced PAECs. DLL4 protein expression was reduced by either *BMPR2* or *JNK1* silencing. Importantly, DLL4 loss was not specific to BMPR2 deficiency and was also seen in our *CAV1* and *SMAD8*/9 LOF models of PAH, and in lung tissue from patients with PAH. This commonality indicates that suppression of DLL4/NOTCH1 signaling might be a central mechanism in PAH. In further support of this, DLL4 was shown to activate NOTCH1/N1ICD signaling and block AKT activation, while reducing cell proliferation and EndoMT in *BMPR2*-silenced PAECs or IPAH ECs. This normalization of endothelial function occurred even in the presence of PAH-associated genetic deficiencies.

DLL4 activates NOTCH1 signaling by liberating N1ICD from the NOTCH1 receptor. N1ICD does not directly bind to DNA, but instead interacts with the transcription factor RBPJ to regulate the expression of NOTCH target genes (7). Notably, two putative RBPJ binding motifs were identified in the *BMPR2* promoter. Exogenous DLL4 increased *BMPR2* mRNA in PAECs from both failed donor controls and IPAH patients. In addition, the overexpression of DLL4 in PAECs activated a reporter gene driven by the *BMPR2* proximal promoter. These findings suggest that BMPR2 deficiency, the most common heritable defect associated with PAH, could be exacerbated or caused by loss or inhibition of DLL4/NOTCH1 signaling.

DLL4/NOTCH1/N1ICD was also found to interact with and target PPARγ, a BMPR2 downstream effector (19) that has anti-inflammatory and anti-proliferative functions in the vasculature (46) and has been described to guard against the development of PAH (18). Reduced PPARγ expression was demonstrated in the lung tissue of patients with PAH and implicated in the proliferative, anti-apoptotic behavior of PAECs from these patients (20). Mice with targeted endothelial (47) or smooth muscle cell (19) deletion of *Pparγ* spontaneously developed PAH with increased right ventricular hypertrophy and muscularization of the distal pulmonary arteries, whereas restoration of PPARγ has been reported to reverse experimental PH (48). Here, DLL4 restored PPARγ levels in ECs isolated from patients with IPAH and also increased *PPARG* mRNA and PPARγ target gene expression in *BMPR2*-silenced PAECs as well as in ECs from IPAH patients. Importantly, DLL4/NOTCH1 signaling-induced PPARγ expression likely inhibits BMPR2 loss-induced AKT activation. Supporting this notion, PPARγ overexpression suppressed AKT activation in *BMPR2*-silenced PAECs. These results indicate that loss of DLL4/NOTCH1 could result in the development of PAH through increased AKT activation along with reduced BMPR2 and PPARγ expression. Consistent with this model of disease pathogenesis, rapamycin (49) and an arginase inhibitor (50) have been shown to attenuate PAH in part by inhibiting AKT activity. In the lung tissue of monocrotaline-treated rats, PPARγ blocked AKT phosphorylation at S473 through the upregulation of PTEN (51), a dual protein/lipid phosphatase that mainly targets PI3K/AKT pathway. The underling mechanism by which PPARγ inhibits AKT activation in the present study warrants further investigation.

Inhibition of the PI3Kα isoform was found to reverse PH in a mouse model of chronic hypoxia, the sugen/hypoxia and MCT-induced rat model by effectively decreasing growth factor induced phosphorylation of AKT in vascular smooth muscle cells (52). However, PI3Kα signaling pathway has been demonstrated to have a cardioprotective role as chronic inhibition of PI3Kα accelerated the progression of heart failure (53). Given that inflammation and excessive PI3K/AKT signaling are common features of PAH and inflammatory cytokines like TNFα can specifically induce the expression of PI3Kδ in ECs, we targeted the PI3Kδ isoform. Leniolisib is an oral PI3Kδ inhibitor, well-tolerated, and recently FDA approved for the treatment of activated PI3Kδ syndrome caused by a gain-of-function mutation in the PI3Kδ gene (PIK3CD) (35). Herein, leniolisib effectively blocked AKT activation in *BMPR2*-silenced PAECs and partially normalized cell proliferation, EndoMT and apoptosis resistance, phenotypic features associated with end stage vascular remodeling in PAH.

In conclusion, we have identified extensive crosstalk among DLL4/NOTCH1 and AKT signaling as well as BMPR2 and PPARγ in human PAECs and related these findings to the pathobiology of vascular remodeling in PAH (Figure 8). Furthermore, PPARγ is identified as an important mechanistic link that is regulated by DLL4/NOTCH1 signaling and blocks AKT activation. DLL4/NOTCH1/PPARγ agonists and/or PI3K/AKT antagonists are attractive therapeutic targets for preventing or even possibly reversing progressive pathologic vasculopathy in PAH.

**Figure 8.**
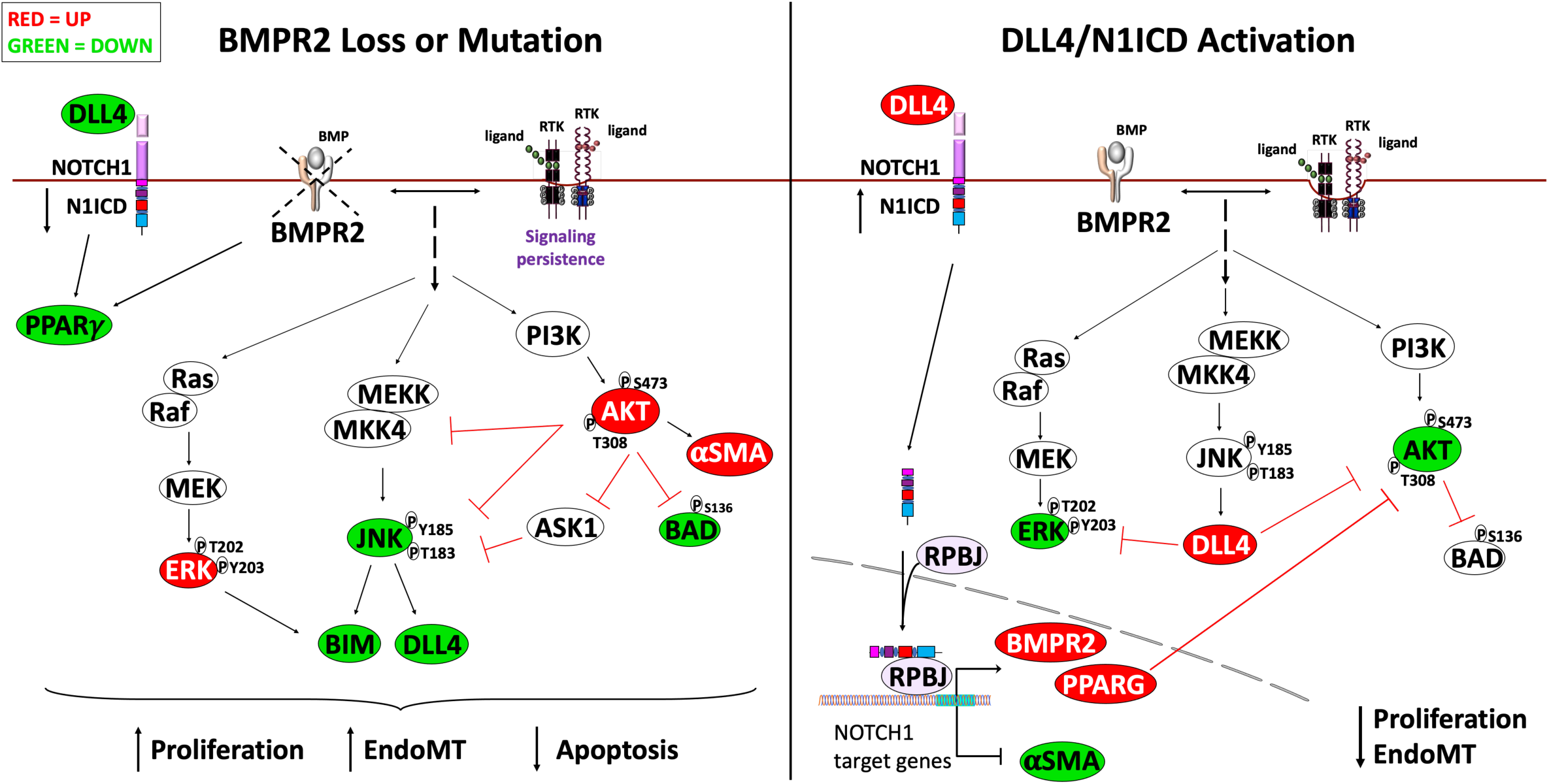
Graphical Abstract. Crosstalk among DLL4/NOTCH1, JNK1, AKT with BMPR2 and PPAR**γ** signaling pathways. BMPR2 loss or Mutation culminates in the hyperactivation of AKT and ERK along with loss of DLL4/NOTCH1, JNK and PPAR**γ** leading to increased proliferation, endothelial to mesenchymal transition (EndoMT) and an apoptosis resistant phenotype. DLL4/N1ICD Activation increases BMPR2/PPAR**γ** expression blocking AKT and ERK activation leading to a decrease in proliferation and EndoMT.

## Acknowledgments

Ling Yi and Vincent Schram at the Microscopy & Imaging Core Facility (National Institute of Child Health and Human Development, NIH) provided invaluable assistance with immunohistochemistry and imaging. Cells and tissue samples used in this study were provided by the Pulmonary Hypertension Breakthrough Initiative (PHBI) Network.

**Author Disclosures** are available with the text of this article.

The opinions expressed in this article are those of the authors and do not represent any position or policy of the NIH, the US Department of Health and Human Services, or the US Government.

## Supplemental Material

Figures E1–E7

Tables E1–E5

## Footnotes

^1^

