## Supplementary material for "Disruption of DLL4/NOTCH1 Causes Dysregulated PPARγ/AKT Signaling in Pulmonary Arterial Hypertension": Online Supplement

### Online Data Supplement

**Supplemental Methods**

*Cell Culturing.* Primary PAECs (Lonza, Frederick, MD) were cultured for a maximum of 6 passages on cell culture flasks coated with type I collagen (50 µg/mL in 0.02 N acetic acid; Corning, Corning, NY) in endothelial basal medium-2 (EBM™-2, Lonza) supplemented with growth factors and 2% serum (EGM™-2 SingleQuot kit, Lonza). Failed donor control (CTRL) and IPAH endothelial cells (ECs) obtained from the Pulmonary Hypertension Breakthrough Initiative (PHBI) (Table E3 donor information) were plated on 0.1 % gelatin (Sigma). Immobilized DLL1 has been reported to mimic cell-to-cell interactions and activate NOTCH signaling (1). Therefore, the same approach was adopted to investigate DLL4 signaling and its ability to activate NOTCH signaling in cultured ECs. In experiments studying exogenous DLL4 effects, cells were seeded on plates pre-coated with recombinant DLL4 (0.5 µg/mL in PBS overnight, R&D Systems; Minneapolis, MN). Cells were maintained at 37°C in a humidified incubator with 5% $CO_2$ and 95% air.

*Gene Silencing.* Primary PAECs were transfected with gene-specific siRNA pools targeting *BMPR2, JNK1, CAV1 or SMAD9* at a final concentration of 10 nM using DharmaFECT-1 (Dharmacon; Lafayette, CO) in Opti-MEM (Invitrogen; Grand Island, NY) for 6 h followed by growth in EGM™-2 for 48 h. Non-targeting siRNA Pool-1 (siGENOME, Dharmacon) was used as a control.

*Immunofluorescence.*Primary PAECs were seeded onto collagen coated 8-well chamber slides (Sigma) at a density of 60,000 cells/well overnight, transfected with *BMPR2*, *CAV1* or control siRNA for 48 h and treated with vehicle (DMSO) or leniolisib (Novartis) for 24 h. Failed donor control or IPAH ECs (50,000 cells/well) were seeded on either BSA or DLL4-coated 8-well chamber slides for 48 h. Cells were then washed 3x with basal medium, fixed with 4% formaldehyde (Polysciences, Inc.) for 20 min at RT, blocked with goat serum (Sigma) plus 0.3% Triton X-100 for 45 min at RT and then washed 3x with PBS supplemented with 0.1% BSA before incubating overnight with primary antibodies. The next day, the slides were washed 3x as above, incubated with secondary antibodies for 1 h, washed 3x as above and mounted using mounting medium with DAPI (4′,6-diamidino-2-phenylindole; Sigma). Cells were imaged using epifluorescence (Leica DMi8 Inverted LED Fluorescence Microscope, Leica, Deerfield, IL) at 20X magnification and analyzed using ImageJ.

*Over-Expression Studies.* Primary PAECs were seeded at 30,000 cells/well on collagen-coated 24-well plates for 48 h followed by 24 h transfection with expression vectors for DLL4 (DLL4/pCMV6-myc-DDK) or empty plasmid (pCMV6-myc-DDK, Origene) along with a *BMPR2* promoter or PPAR-luciferase reporter and pGL4.74 [hRluc/TK, *Renilla*-luciferase] plasmid (Promega, Madison, WI) using Lipofectamine$^{TM}$ Stem reagent (ThermoFisher). The *BMPR2* promoter was designed by cloning the -1200/+100 human *BMPR2* promoter into the pGL4.15[luc2p/Hygro] vector (Promega). The PPAR reporter, containing 3 copies of PPAR response element upstream of the reporter gene 3-thymidine kinase-luciferase (PPAR-LUC), was kindly provided by Dr. Ronald M. Evans (2). PAECs were also seeded on either BSA or DLL4 coated plates for 48 h then transfected with the *BMPR2* promoter-luciferase reporter for an additional 24 h. Lysates were harvested using 1x passive lysis buffer (Promega) and analyzed using the Dual-Luciferase reporter assay system (Promega). Data was normalized to Renilla-luciferase from the same sample.

For protein lysates, PAECs were plated on either BSA or DLL4 overnight followed by *BMPR2* silencing for 24 h. The cells were then transfected with either empty plasmid (CMV) or PPARγ (PPARγ/ pCMV6-myc-DDK) for 24 h before adding RIPA buffer (ThermoFisher Scientific) supplemented with protease and phosphatase inhibitor cocktail (ThermoFisher Scientific). Additionally, protein lysates were also collected 24 h after over-expressing empty plasmid or DLL4.

*RNA Isolation, cDNA Synthesis and Quantitative Real-Time PCR.* Total RNA was extracted using RNeasy Mini Kit (Qiagen, Valencia, CA) according to the manufacturer's instructions, including DNase I treatment. The iScript cDNA Synthesis Kit (Bio-Rad; Hercules, CA) was used to synthesize cDNA and quantitative real-time PCR (qRT-PCR) analysis was performed using iTaq Universal SYBR Green Supermix with ROX (Bio-Rad) on a Biosystems ViiA$^{TM}$7 instrument. Gene expression was normalized to β-actin and delta cycle thresholds were tested for significance. Primer sequences of target genes are listed in Table E4.

*Western Blot.* Whole cell protein lysates from PAECs, unused failed donor and IPAH cells were lysed with RIPA buffer supplemented with protease and phosphatase inhibitor cocktail. Protein lysates (30 μg) fractionated by 4-12% SDS-PAGE were transferred on nitrocellulose membranes, blocked at RT for 1 h and incubated overnight at 4° C with primary antibody (Table E5). Blots were imaged with ChemiDoc MP Imaging System (BioRad) and densitometry analysis performed with Image Lab software (version 6.1; BioRad).

*Cell Proliferation Assays.* Primary PAECs were transfected with siCTRL or siBMPR2 for 48 h in T75 flasks, detached with 0.5% trypsin-EDTA and re-seeded (5000 cells/well) on BSA or DLL4 coated 96-well plates. Likewise, failed donor control or IPAH ECs were seeded on BSA or DLL4 coated 96-well plates and cell proliferation over 72 h was quantitated by incorporation of 5-bromo-2-deoxyuridine (BrdU) using a chemiluminescence-based ELISA kit (Sigma).

*Cell Apoptosis.* Primary PAECs transfected with either BMPR2 or control siRNA for 48 h were replated on collagen-coated 96-well plates at a density of 5000 cell/well overnight. The following day, media was replaced with either complete media or media without serum or growth factors for 24 h. An equal volume of Caspase-Glo 3/7 reagent (Promega) was added to each well for 30 min and luminescence was read with a GloMax® plate reader (Promega). For flow cytometry, primary PAECs were plated on collagen-coated T25 flasks at a density of 220,000 cells/flask. The next day, cells were transfected with either control, *BMPR2*, *JNK1* or *CAV1* siRNA for 48 h. Cell media was then replaced with either complete media or media without serum or growth factors and the addition of either vehicle (DMSO) or leniolisib for another 24 h. Cells were harvested, stained with annexin V and propidium iodide (PI) (BD Pharmingen™) and evaluated using either a MACSquant® (Miltenyi) or BD LSRFortessa™ (BD Biosciences) analyzer. Data files were then uploaded and analyzed using FlowJo software version 10.4.2 (Treestar, Ashland, OR).

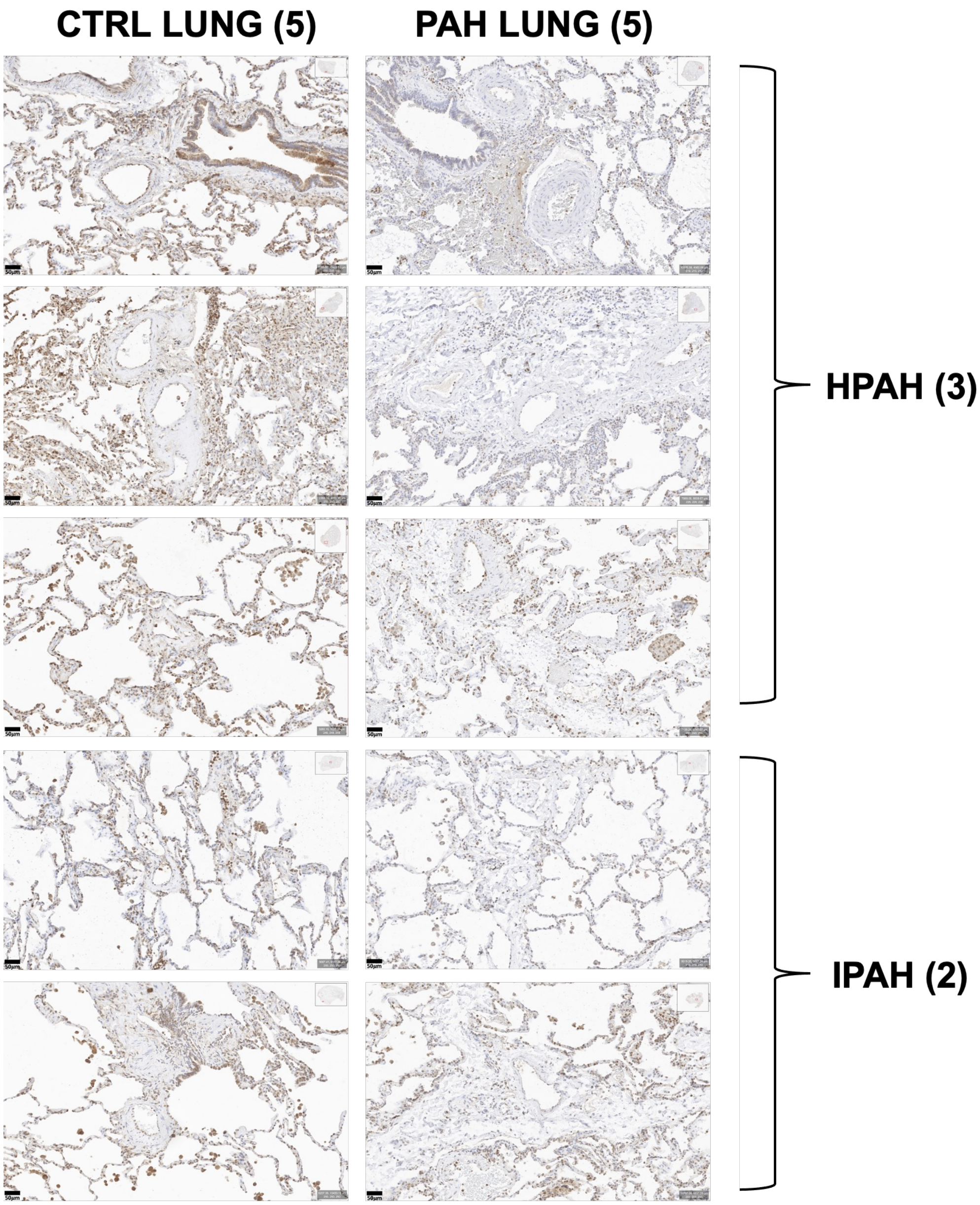


**Figure E1**. **DLL4 protein is reduced in lung of patients with PAH.** Immunohistochemical staining of DLL4 in paraffin embedded lung of failed donor controls (CTRL; n = 5), HPAH (n = 3) and IPAH (n = 2). Scale bar, 50µm.

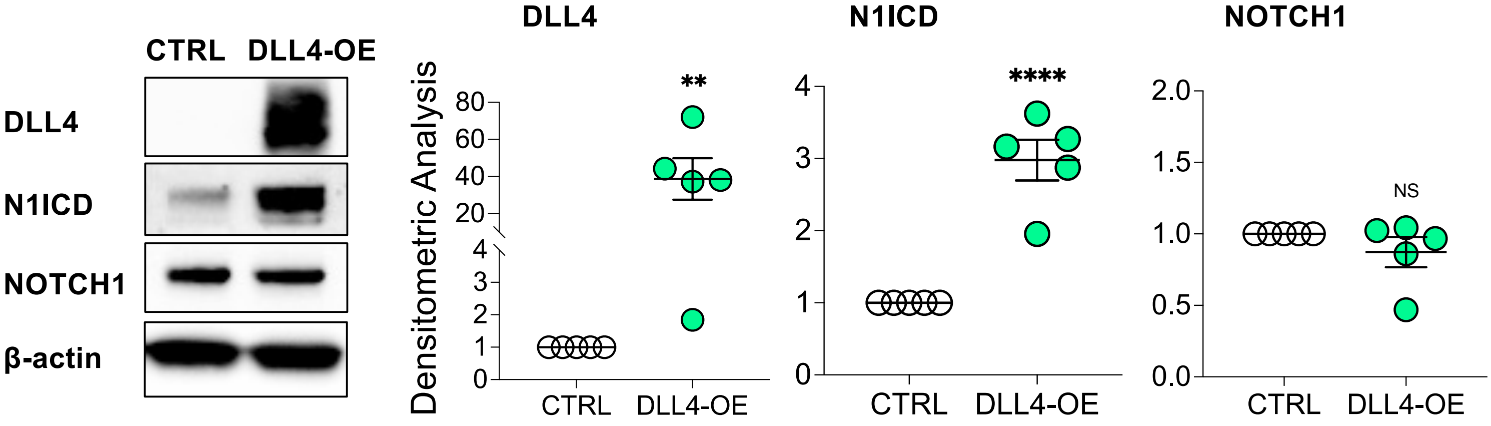


**Figure E2. DLL4 overexpression increases N1ICD expression in primary human pulmonary artery endothelial cells (PAECs).** PAECs were transfected with empty vector (CTRL) or DLL4 overexpression plasmid (DLL4-OE) for 24 h and total protein lysates were collected. Representative Western blots are shown for DLL4, N1ICD and NOTCH1 (n=5). Densitometric analysis of each protein is relative to β-actin and normalized to its corresponding control. Data presented as mean±SEM; paired t-test: ***P* < 0.01; *****P* < 0.001; NS, not significant.

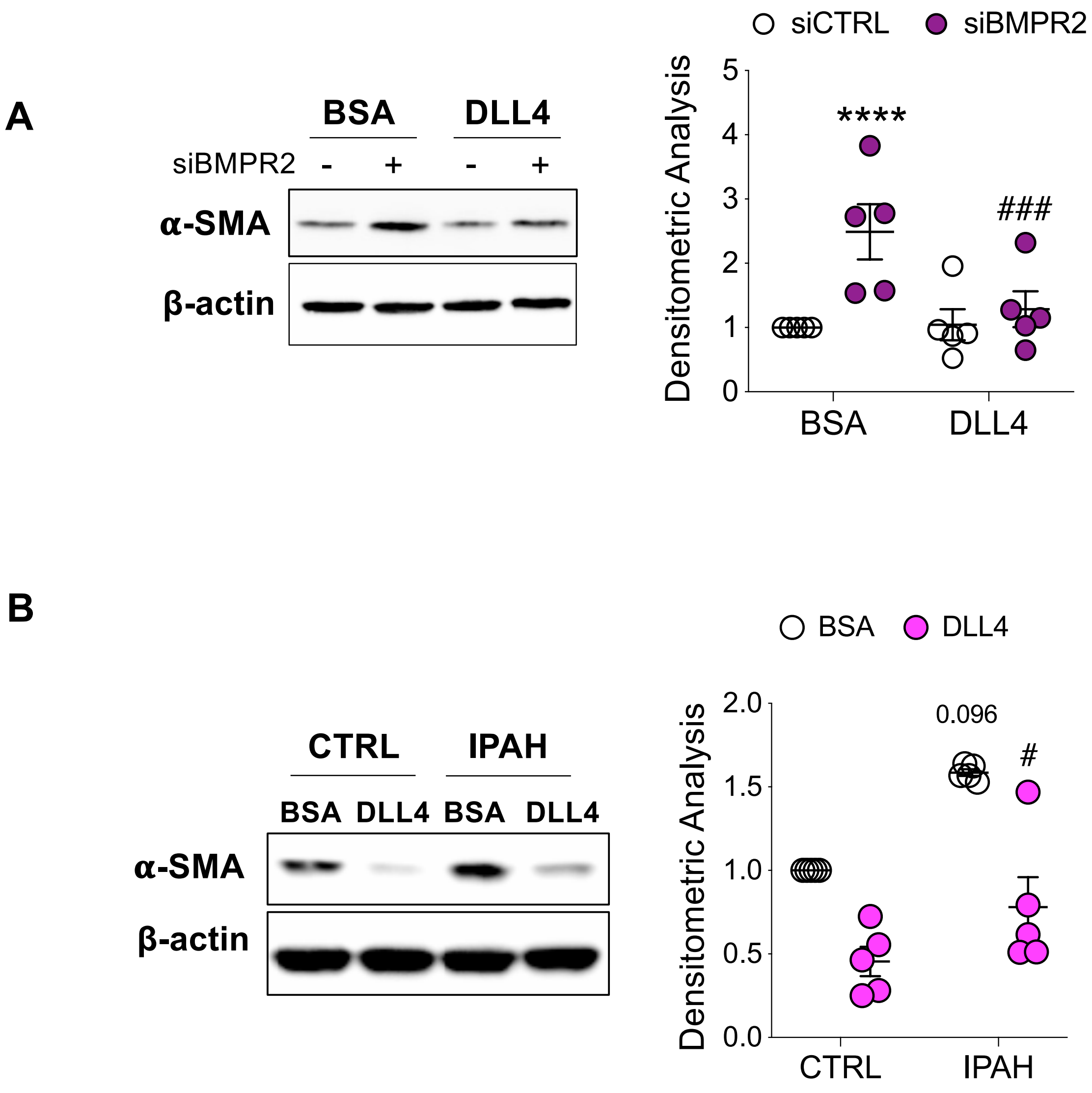


**Figure E3. Exogenous DLL4 blocks α-SMA (ACTA2) induction in *BMPR2*-silenced human primary pulmonary artery endothelial cells (PAECs) and in PAECs from patients with idiopathic pulmonary arterial hypertension (IPAH).** (A) PAECs were grown on BSA or DLL4 coated plates and transfected with control (siCTRL) or BMPR2 (siBMPR2) siRNA for 48 h (n=5). (B) PAECs from failed donors (CTRL) and IPAH patients were grown on either BSA or DLL4 coated plates for 48 h (n=5). For both **(A)** and **(B)** total protein lysates were collected, and representative Western blots of α-SMA and β-actin are shown. Densitometric analyses of each protein relative to β-actin were normalized to its corresponding control. Data presented as mean±SEM; 2-way ANOVA with Tukey HSD: (A) $^{****}P < 0.001$ (siCTRL-BSA versus siBMPR2-BSA); $^{###}P < 0.005$ (siBMPR2-BSA versus siBMPR2-DLL4); (B) $P = 0.09$ (CTRL-BSA versus IPAH-BSA); $^{\#}P < 0.05$ (IPAH-BSA versus IPAH-DLL4).

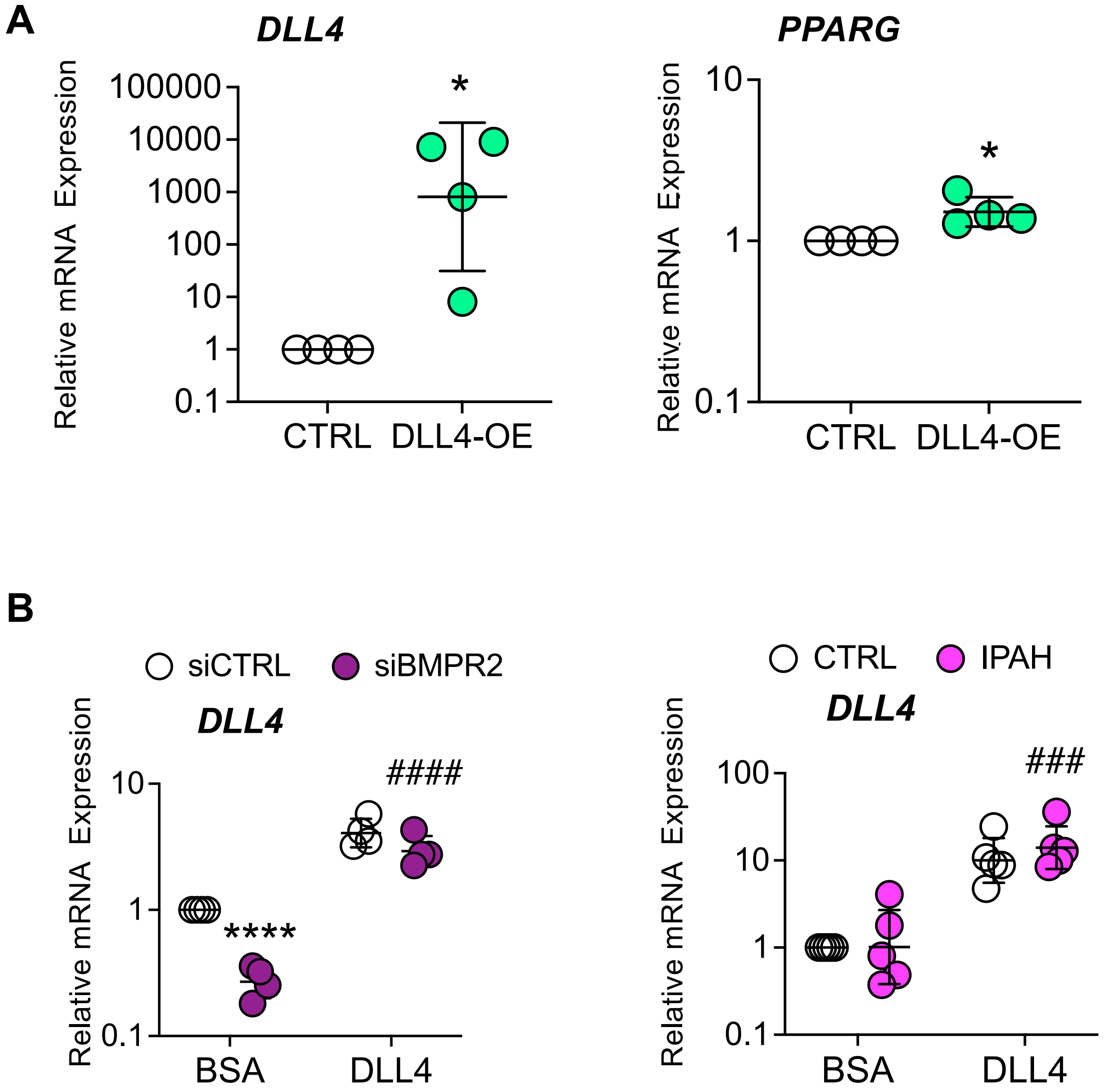


**Figure E4. DLL4 overexpression or immobilized DLL4 increases DLL4 and PPARG mRNA in human primary pulmonary artery endothelial cells (PAECs) or cells from patients with idiopathic pulmonary arterial hypertension (IPAH).** (A) Quantitative RT-PCR of DLL4 and PPARG mRNA in human primary pulmonary artery endothelial cells (PAECs) transfected with either empty vector (CTRL) or DLL4 overexpression plasmid (DLL4-OE) for 24 h (n=3). (B) Quantitative RT-PCR of DLL4 in PAECs transfected with control (siCTRL) or BMPR2 (siBMPR2) siRNA, or PAECs from failed donors (CTRL) and IPAH patients grown on BSA or DLL4 coated plates (n=5). Data are presented as the geometric mean ± SD. (A) Paired t-test. (B) 2-way ANOVA with Tukey HSD: ****P < 0.001 (siCTRL-BSA versus siBMPR2-BSA); ####P < 0.001 (siBMPR2-BSA versus siBMPR2-DLL4); ###P < 0.005 (IPAH-BSA versus IPAH-DLL4).

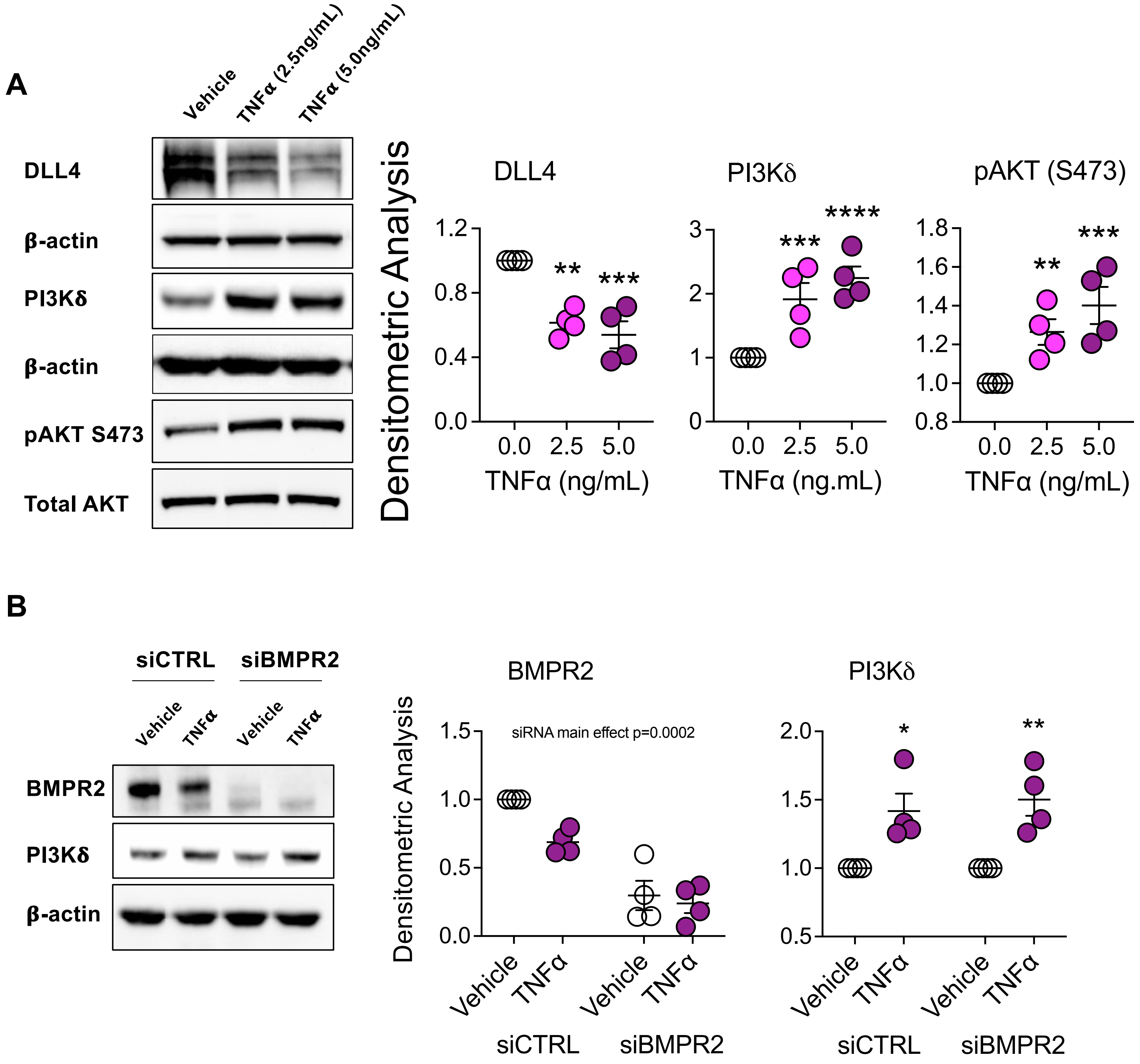


**Figure E5. TNFα suppresses DLL4 expression, induces PI3Kδ and activates AKT in human primary pulmonary artery endothelial cells (PAECs).** (A) PAECs were treated with vehicle or TNFα at the indicated concentrations for 24 h. Total protein lysates were then collected and analyzed by Western blotting for expression of DLL4, PI3Kδ, and phosphorylated AKT (pAKT S473) or p50 NFκB (n=4). (B) PAECs were transfected with control (siCTRL) or BMPR2 (siBMPR2) siRNA for 48 h and then exposed to TNFα (5 ng/mL) for 24 h. Cytoplasmic fractions were prepared and analyzed by Western blotting for expression of BMPR2 and PI3Kδ (n=4). Densitometric analysis relative to β-actin or total AKT was normalized to its corresponding control. Representative Western blots are shown. Data are presented as mean±SEM. (A) 1-way ANOVA with Tukey HSD; ***P* < 0.01; ****P* < 0.05; *****P* < 0.001 (vehicle vs treatment); (B) 2-way ANOVA with Tukey HSD in B: **P* < 0.05; ***P* < 0.01 (vehicle vs treatment).

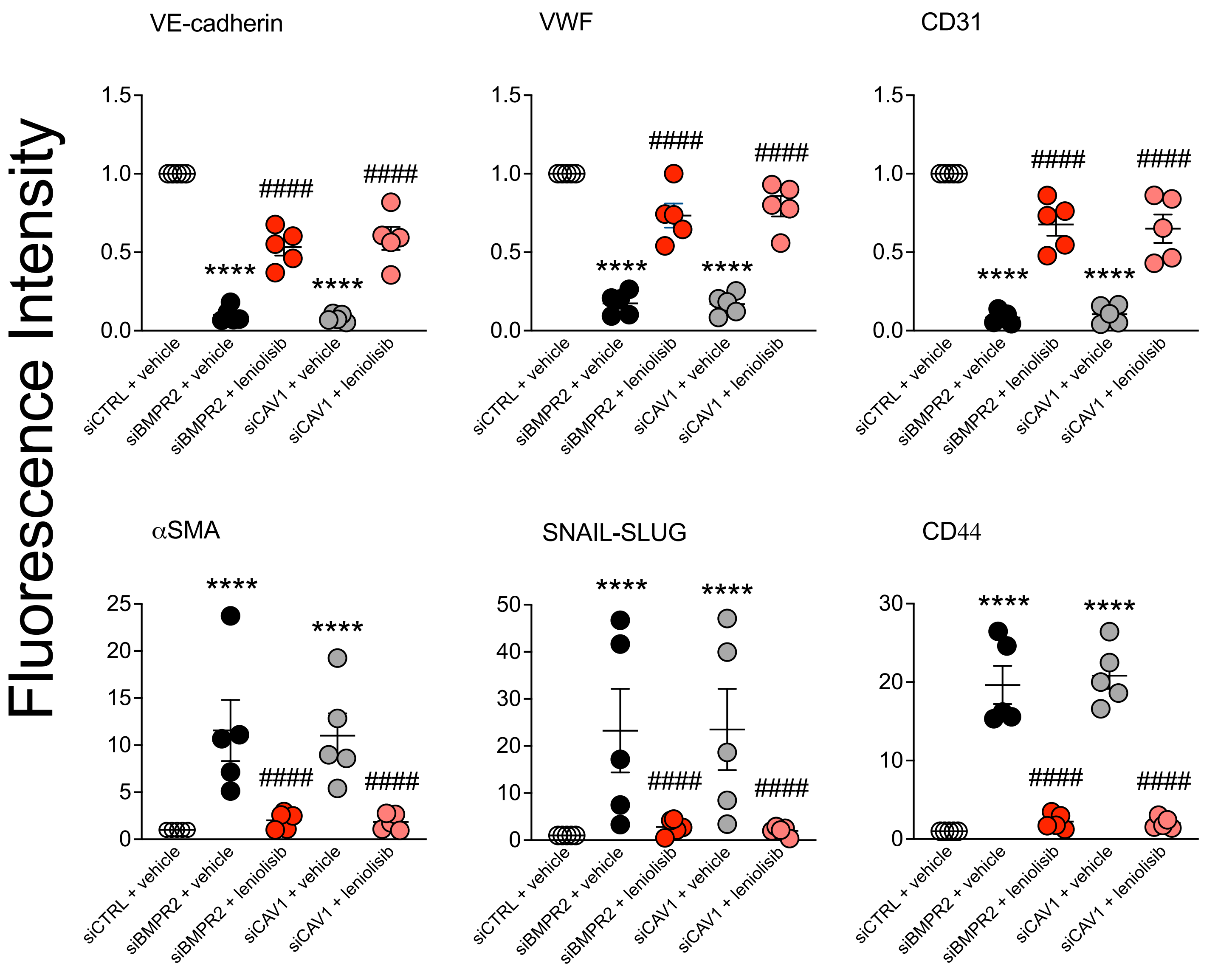


**Figure E6. Leniolisib inhibits endothelial-mesenchymal transition (EndoMT) in *BMPR2* and *CAV1* silenced human primary pulmonary artery endothelial cells (PAECs).** Fluorescence intensity of endothelial markers (VE-cadherin, VWF and CD31) and mesenchymal markers (α-SMA, SNAIL/SLUG and CD44) in PAECs transfected with control (siCTRL), BMPR2 (siBMPR2), or CAV1 (siCAV1) siRNA for 48 h. BMPR2- or CAV1-silenced cells were then incubated for an additional 24 hours without or with 10 μM leniolisib (n=5). Data are presented as mean±SEM; 2-way ANOVA with Tukey HSD: ****$P$ < 0.001 (siCTRL-vehicle versus siBMPR2-vehicle or siCTRL-vehicle versus siCAV1-vehicle); ####$P$ < 0.001 (siBMPR2-vehicle versus siBMPR2-leniolisib or siCAV1-vehicle versus siCAV1-leniolisib).

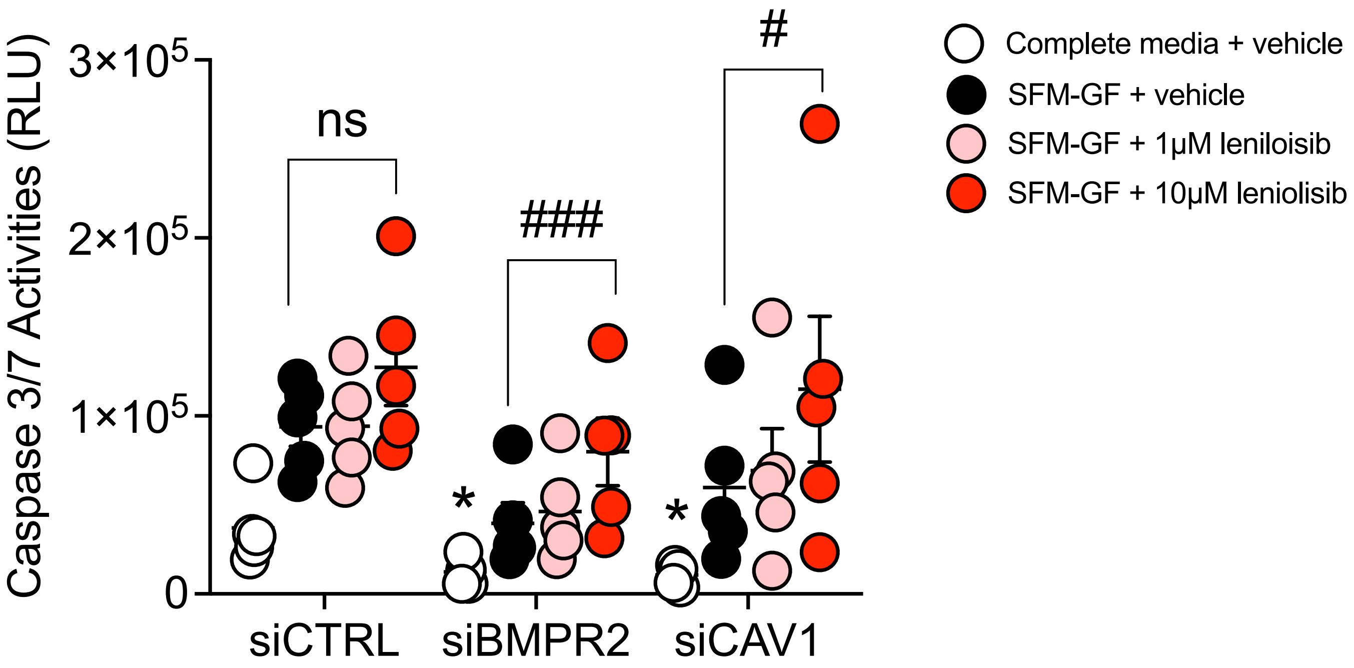


**Figure E7. Leniolisib induces caspase 3/7 activity, a marker of apoptosis, in BMPR2- and CAV1-silenced human primary pulmonary artery endothelial cells (PAECs).** PAECs were transfected with control (siCTRL), BMPR2 (siBMPR2) or CAV1 (siCAV1) siRNA for 48 h then replated and treated with vehicle (DMSO), serum free media minus growth factors (SFM-GF) plus vehicle, SFM-GF plus 1 μM leniolisib or SFM-GF plus 10 μM leniolisib for an additional 24 h (n=5). Data are presented as mean±SEM; 2-way ANOVA with Tukey HSD: *$P < 0.05$ (siCTRL-complete media + vehicle versus siBMPR2-complete media + vehicle or siCTRL-complete media + vehicle versus siCAV1-complete media + vehicle); ###$P < 0.005$ (siBMPR2-SFM-GF + vehicle versus siBMPR2-SFM-GF + 10 μM leniolisib); #$P < 0.05$ (siCAV1-SFM-GF + vehicle versus siCAV1-SFM-GF + 10 μM leniolisib).

#### Table E1: Patients' characteristics; Frozen Tissue

| PHBI # | Age | Race | Gender | Diagnosis | BMPR2 Mutation |
|---|---|---|---|---|---|
| AH-022 | 57 | White | Female | Failed Donor | No |
| AH-028 | 43 | White | Female | Failed Donor | No |
| BA-037 | 60 | White | Female | Failed Donor | No |
| BA-046 | 55 | White | Female | Failed Donor | No |
| BA-054 | 33 | Hispanic | Female | Failed Donor | No |
| BA-059 | 34 | American Asian | Female | Failed Donor | No |
| UA-018 | 50 | White | Female | Failed Donor | No |
| UA-020 | 51 | Hispanic | Female | Failed Donor | No |
| CC-030 | 63 | White | Female | IPAH | No |
| ST-001 | 10 | Asian/Pacific Islander | Female | IPAH | No |
| ST-037 | 39 | White | Female | IPAH | No |
| VA-014 | 33 | White | Female | IPAH | Yes |
| CC-015 | 32 | White | Female | HPAH | Yes |
| CC-032 | 54 | White | Female | HPAH | Yes |
| UA-023 | 23 | White | Female | HPAH | Yes |
| VA-010 | 33 | Black | Female | HPAH | Yes |

#### Table E2: Patients’ characteristics for Immunohistochemistry

| PHBI # | Age | Race | Gender | Diagnosis | BMPR2 Mutation |
|---|---|---|---|---|---|
| UA-018 | 50 | White | Female | Failed Donor | No |
| UA-021 | 49 | White | Female | Failed Donor | No |
| AH-022 | 57 | White | Female | Failed Donor | No |
| BA-046 | 55 | White | Female | Failed Donor | No |
| BA-059 | 34 | American Asian | Female | Failed Donor | No |
| ST-012 | 26 | American Asian | Female | IPAH | Yes |
| VA-014 | 33 | White | Female | IPAH | Yes |
| VA-010 | 33 | Black | Female | HPAH | Yes |
| UA-023 | 23 | White | Female | HPAH | Yes |
| CC-033 | 25 | White | Female | HPAH | Yes |

#### Table E3: PHBI Subject Information

| PHBI # | Age | Race | Gender | Diagnosis | Materials |
|---|---|---|---|---|---|
| UA-015 | 36 | White | Female | Failed Donor | LMVEC |
| BA-037 | 60 | White | Female | Failed Donor | LMVEC |
| BA-046 | 55 | White | Female | Failed Donor | LMVEC |
| AH-022 | 57 | White | Female | Failed Donor | LMVEC |
| BA-061 | 46 | White | Female | Failed Donor | LMVEC |
| UA-018 | 50 | White | Female | Failed Donor | LMVEC |
| CC-013 | 27 | White | Female | IPAH | LMVEC |
| VA-011 | 32 | White | Female | IPAH | LMVEC |
| ST-040 | 16 | White | Female | IPAH | LMVEC |
| VA-017 | 40 | White | Female | IPAH | LMVEC |
| ST-028 | 40 | White | Female | IPAH | LMVEC |
| UA-026 | 55 | White | Female | IPAH | LMVEC |

#### Table E4. Quantitative real-time PCR primer sequences

| Gene | Forward | Reverse |
|---|---|---|
| BMPR2 | 5′-CACTGCGGCTGCTTCGCAGA-3′ | 5′-AGCAGGTGCTACCTTTCGAGCA-3′ |
| PPARγ | 5′-GGATTCAGCTGGTCGATATCAC-3′ | 5′-GTTTCAGAAATGCCTTGCAGT-3′ |
| FABP4 | 5′-ATCACATCCCCATTCACACT-3′ | 5′-ACTTGTCTCCAGTGAAAACTTTG-3′ |
| CYP1A1 | 5′-TGGAGATTGGGAAAAGCATGA-3′ | 5′-GAACCTTCCCTGATCCTTGTG-3′ |
| PGK1 | 5′-GACAGCAGCCTTAATCCTCTG-3′ | 5′-CTAACAAGCTGACGCTGGA-3′ |
| HK1 | 5′-TCCCAACAATGAGTCCAACC-3′ | 5′-GCCACGATGTAGTCACCTTAC-3′ |
| β-ACTIN | 5′-CCGCCGCCAGCTCACCAT -3′ | 5′-ACCCATGCCCACCATCAGGC-3′ |
| DLL4 | MIQE CONTEXT SEQUENCE:<br>TGAGCAAACCAGCACCCTCACAAGG<br>CTGCGCTACTCTTACCGGGTCATCT<br>GCAGTGACAACTACTATGGAGACAA<br>CTGCTCCCGCCTGTGCAAGAAGCGC<br>AATGACCACTTCGGCCAC | |

### Table E5

| REAGENT or RESOURCE | SOURCE | IDENTIFIER |
|---|---|---|
| **Antibodies and Reagents** | | |
| BMPR2 | BD Biosciences | 612292, RRID:AB_399609 |
| pAKT S473 | Cell Signaling | 9271S, RRID:AB_329825 |
| pAKT T308 | Cell Signaling | 9275S, RRID:AB_329828 |
| Total AKT | Cell Signaling | 9272S, RRID:AB_329827 |
| pERK | Cell Signaling | 9101S, RRID:AB_331646 |
| Total ERK | Cell Signaling | 9102S, RRID:AB_330744 |
| pJNK | Cell Signaling | 4671S, RRID:AB_331338 |
| Total JNK | Cell Signaling | 9258S, RRID:AB_2141027 |
| Cleaved CASPASE 3 | Cell Signaling | 9661S, RRID:AB_2341188 |
| BCL2L11 (BIM) | Abcam | Ab7888, RRID:AB_2065161 |
| NOTCH1, cleaved (Val1744) (N1ICD) | Cell Signaling | 4147S, RRID:AB_2153348 |
| NOTCH1 | Cell Signaling | 3608S, RRID:AB_2153354 |
| NOTCH2 | Cell Signaling | 5732S, RRID:AB_10693319 |
| N4ICD | Cell Signaling | 2423S, RRID:AB_2151366 |
| DLL4 | Cell Signaling | 96406S, RRID:AB_2800263 |
| pBAD S136 | Cell Signaling | 4366S, RRID:AB_10547878 |
| Total BAD | Cell Signaling | 9292S, RRID:AB_331419 |
| PPARγ | Cell Signaling | 2443S, RRID:AB_823598 |
| β-Actin (ACTB) | Sigma | A-3854, RRID:AB_262011 |
| α-Smooth Muscle Actin (ACTA2) | Sigma | A-5228, RRID:AB_262054 |
| CD31 (immunofluorescence) | BD BioSciences | 555445, RRID:AB_395838 |
| CD31 (immunofluorescence) | Agilent | M0823, RRID:AB_2114471 |
| CDH5 (VE-cadherin; immunofluorescence) | Cell Signaling | 2518S, RRID:AB_2077970 |

| | | |
|---|---|---|
| VWF (immunofluorescence) | Abcam | ab154193 |
| VEGFR2 (immunofluorescence) | Cell Signaling | 2479S, RRID:AB_10698606 |
| CD44 (immunofluorescence) | Abcam | ab189524, RRID:AB_2885107 |
| SNAI1/SLUG (immunofluorescence) | Abcam | ab167609 |
| Horseradish peroxidase-conjugated goat anti-rabbit | Bio-Rad | 170-6515, RRID:AB_11125142 |
| Horseradish peroxidase-conjugated goat anti-mouse | Bio-Rad | 170-6516, RRID:AB_11125547 |
| Donkey anti-Mouse IgG (H+L) Cross-Adsorbed secondary, Alexa Fluor 488 | ThermoFisher Scientific | A-21202, RRID:AB_141607 |
| Donkey anti-Rabbit IgG (H+L) Cross-Adsorbed secondary, Alexa Fluor 647 | ThermoFisher Scientific | A-32795, RRID:AB_2762835 |
| Prolong Diamond Antifade Mountant | ThermoFisher Scientific | P36961 |
| **Recombinant proteins** | | |
| Recombinant Human DLL4 protein | R&D Systems | 1506-D4-050 |
| **Experimental models** | | |
| Human pulmonary artery endothelial cells (HPAEC) | Lonza | CC-2530 |
| IPAH, see Table S1 | PHBI Penn Cell Center | 02544 |
| Non-Diseased (Controls), see Table S1 | PHBI Penn Cell Center | 02544 |
| **Cell Culture** | | |
| EGM-2 BulletKit (CC-3156 & CC-4176) | Lonza | CC-3162 |
| EGM-2 MV BulletKit (CC-3156 & CC-4177) | Lonza | CC-3202 |
| Opti-MEM | Gibco | 31985077 |
| **Critical commercial assays** | | |
| Caspase-Glo 3/7 | Promega | G8093 |

| | | |
|---|---|---|
| FITC Annexin V Apoptosis Detection Kit | BD Pharmingen | 556547 |
| Cell Proliferation ELISA, BrdU (chemiluminescent) | Sigma | 11669915001 |
| NE-PER Nuclear and Cytoplasmic Extraction kit | ThermoFisher Scientific | 78835 |
| Lipofectamine™ Stem reagent | ThermoFisher Scientific | STEM00001 |
| iScript™ cDNA Synthesis Kit | Bio-Rad | 170-8891 |
| iTaq™ Universal SYBR® Green Supermix | Bio-Rad | 1725124 |
| **Oligonucleotides** | | |
| Primers for qPCR, see Table S2 | | |
| **Recombinant DNA** | | |
| Vector PPRE3-TK-LUC; 3 copies of PPRE upstream of the thymidine kinase promoter fused to the luciferase gene | | Kindly provided by Ronald M. Evans |
| pGL4[luc2P/RBP-Jk-RE/Hygro] | Promega | CS173601 |
| pGL4.15[luc2P/BMPR2/Hygro] | Promega | custom |
| pRL-TK Renilla luciferase | Promega | custom |
| PPAR$\gamma$ expression plasmid | Addgene | 8895 |
| DLL4 expression plasmid | Origene | custom |
